# Intrinsic/extrinsic duality of large-scale neural functional integration in the human brain

**DOI:** 10.1101/2020.04.21.053579

**Authors:** Martin Sjøgård, Mathieu Bourguignon, Lars Costers, Alexandru Dumitrescu, Tim Coolen, Liliia Roshchupkina, Florian Destoky, Julie Bertels, Maxime Niesen, Marc Vander Ghinst, Jeroen van Schependom, Guy Nagels, Charline Urbain, Philippe Peigneux, Serge Goldman, Mark W. Woolrich, Xavier De Tiège, Vincent Wens

## Abstract

Human brain activity is not merely responsive to environmental context but includes intrinsic dynamics, as suggested by the discovery of functionally meaningful neural networks at rest, i.e., even without explicit engagement of the corresponding function. Yet, the neurophysiological coupling mechanisms distinguishing intrinsic (i.e., task-invariant) from extrinsic (i.e., task-dependent) brain networks remain indeterminate. Here, we investigated functional brain integration using magnetoencephalography throughout rest and various tasks recruiting different functional systems and modulating perceptual/cognitive loads. We demonstrated that two distinct modes of neural communication continually operate in parallel: extrinsic coupling supported by phase synchronization and intrinsic integration encoded in amplitude correlation. Intrinsic integration also contributes to phase synchronization, especially over short (second-long) timescales, through modulatory effects of amplitude correlation. Our study establishes the foundations of a novel conceptual framework for human brain function that fundamentally relies on electrophysiological features of functional integration. This framework blurs the boundary between resting-state and task-related neuroimaging.

## Introduction

Human brain functions emerge from neural computations coordinated across distant areas of the nervous system. This process of functional integration is partly structured by the underlying neuroanatomical connectivity but differs from it by its functional nature, i.e., brain networks are generated dynamically according to stimulus- or task-related demands. Functional integration was thus first considered as an *extrinsic*, contextdependent property^1^ of brain activity enabling the recruitment of the network configuration required by current environmental demands. However, neuroimaging studies also identified several resting-state networks (RSNs)^2,3^, which anatomically resemble extrinsic networks but spontaneously emerge at rest, i.e., in the absence of any explicit stimulation or task.^4^ This discovery fundamentally shifted the conception of brain functional organization. Besides reacting to environmental events, the brain continually probes network configurations prospectively, presumably to favor the efficient recruitment of extrinsic networks when needed.^3^ This hypothesis is suggestive of an *intrinsic* mode of functional integration not directly driven by environmental demands.^5,6^ Intrinsic functional integration has hitherto been mostly studied through resting-state connectivity. Still, this approach is insufficient to discriminate truly intrinsic (i.e., contextually invariant^1^) from extrinsic (i.e., including at least some context-dependent features^1^) functional systems. Indeed, resting-state activity includes spontaneous cognitive processes (e.g., mind wandering), which are extrinsic as they are subjected to cognitive and perceptual changes.^7^ Disclosing intrinsic integration in the sense of being stimulation/task independent would rather reveal purely mechanistic, endogenous brain dynamics^8^ maintaining, e.g., synaptic homeostasis^5^. Yet, empirical disambiguation of intrinsic/extrinsic neural features is still lacking, despite their critical significance for brain organization. Accordingly, fundamental questions remain open. Are RSNs intrinsic or do they exhibit extrinsic properties? How can intrinsic and extrinsic neural communication coexist? Do they rely on different neurophysiological mechanisms, and are they functionally inter-related?

We addressed these questions by considering distinct electrophysiological markers of functional connectivity, i.e., measures of statistical inter-dependence reflecting aspects of neural communication within the brain^9^. Connectivity measures between the oscillatory activity of neural assemblies can be broadly separated into amplitude and phase coupling,^10^ which provide non-redundant information about brain interactions.^11^ Both are assumed to contribute to intrinsic networks^6^ since RSNs are identifiable from amplitude correlation^12–14^ and short-lived events of phase synchrony^15^, but these clues are limited to restingstate connectivity and do not establish their stimulation/task independence. The role of phase synchronization in distributed task-evoked neural activity^16,17^ further suggests that it supports extrinsic coupling. We recorded the neuroelectric activity of healthy adults with magnetoencephalography (MEG)^18^, both at rest and during several tasks recruiting different functional systems and varying cognitive/perceptual load. We assessed whether these functional connectivity measures changed accordingly. This experimental design allowed to disambiguate the electrophysiological mechanisms of intrinsic and extrinsic coupling, the former identified by task independence and the latter by task-related modulations. It also allowed us to investigate extrinsic properties of RSNs and the functional relevance of intrinsic integration for extrinsic coupling.

## Results

### Network-level functional connectivity

We mapped the large-scale human connectome by considering all connections between 155 nodes densely parcellating the cortex into RSN areas (Figure 1a) and defined from resting-state functional magnetic resonance imaging (fMRI).^19^ Neuroelectric activity at these nodes was reconstructed using minimum norm estimation, with geometric correction of non-physiological leakage connectivity induced by magnetic field spread across two nodes.^20^ We measured frequency-specific amplitude and phase couplings from 4 to 30 Hz using time-averaged envelope correlation and phase-locking value respectively (Figure 1b).^10^ We also applied power regression to exclude connectivity estimation biases caused by local task-related changes in signal-to-noise ratio^21^. To identify intrinsic/extrinsic features, these functional connectomes were computed separately at rest and during task performance. We probed different functional systems with MEG recordings from experiments conducted in several groups of healthy adults engaged in motor sequence learning (*N* = 26 subjects), speech-in-noise comprehension in a purely auditory (*N* = 25) and an audiovisual (*N* = 25) modality, covert language production (*N* = 30), or working memory (*n*-Back, *N* = 35). In each experiment, perceptual or cognitive loads were modulated by varying task difficulty (i.e., motor sequence complexity, auditory noise level or type, language production process, and working memory load). Figures 1c,d illustrate the broadband connectivity strength within and between RSNs throughout the n-Back experiment (Supplementary Results S1, for the other datasets). Amplitude correlation was systematically strongest within RSNs, both at rest^12–14^ and during task, suggesting that RSNs reflect intrinsic integration. Time-averaged phase coupling was not structured that way, and appeared extrinsic as it changed from rest to task.

**Figure 1:**
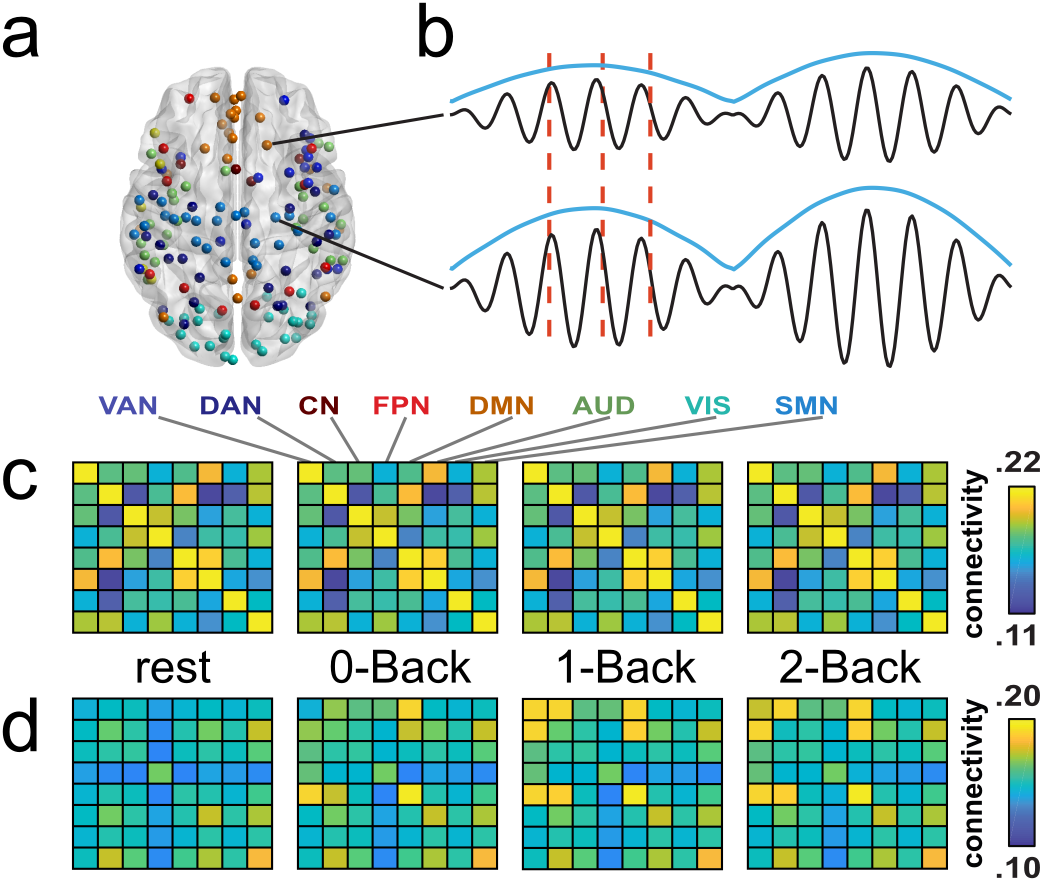
Electrophysiological measures of connectivity. **a**, Network-based brain parcellation used in this study and derived from a meta-analysis of restingstate fMRI.^19^ **b**, Amplitude and phase coupling between reconstructed oscillatory brain signals. Amplitude correlation gauges the covariation of their Hilbert envelope (overhanging blue curves). The phase-locking value evaluates the consistency of the phase delay between them, as measured by the time lag between subsequent local maxima (vertical red dotted lines). Parts **c** and **d** show the broadband (i.e., average across frequency bands), network-level (i.e., average across RSN nodes) connectivity patterns for these two coupling measures throughout the *n*-Back conditions. **c**, Amplitude correlation. **d**, Phase coupling. VAN, ventral attentional network; DAN, dorsal attentional network; CN, control-executive network; FPN, fronto-parietal network; DMN, default-mode network; AUD, auditory network; VIS, visual network; SMN, sensorimotor network.

That said, just because amplitude correlation is weaker between RSNs does not mean that it is absent.^22,23^ Amplitude correlation was significantly above noise level (estimated from empty-room MEG recordings) for 75 - 87% of intra-RSN links and 51 - 62% of cross-RSN links (detection rates at *p* < 0.05 corrected for multiple comparisons over all connections, frequencies, and conditions; range across datasets). Accordingly, the whole connectome exhibits amplitude coupling, also between RSNs. Phase locking significantly above noise level was much scarcer but still observable in all datasets (intra-RSN detection rates: 8 - 11%, cross-RSN: 6 - 10%), including at rest. The coarse RSN connectivity plots of Figure 1c do not provide clear insight into the intrinsic/extrinsic nature of cross-RSN amplitude correlation (because they are dominated by intra-RSN correlations). They might also conceal extrinsic intra-RSN amplitude coupling at finer spatial and spectral scales. Likewise, Figure 1d might miss intrinsic phase coupling (possibly due to its transient nature^15^). We thus turned to a systematic classification of frequency-specific connections in terms of their task dependence. Functional connectivity at finer timescales is considered afterwards.

### Task-dependent states of amplitude correlation

For each dataset and coupling type, we sought to identify the possible ways connectivity was affected by task. We partitioned connections and frequencies in terms of how they vary with task, using *k*-means clustering with robust estimation of the number of different patterns^24^. We excluded beforehand frequency-specific connections not significantly exceeding noise level in any condition of the experiment, as noise could spuriously generate task-independent patterns. This allowed to detect distinct *task-dependent connectivity states*, i.e., functional networks reacting differently to task. Classification of amplitude correlation revealed one or two states per dataset. Figure 2a shows the two *n*-Back states. All these states exhibited a homogeneously high degree of similarity in their task dependence, independently of the RSN structure (Figure 2a, top, for the *n*-Back) and systematically across all experiments (Supplementary Results S2). They were also broadband except for the two *n*-Back states, which split the connectivity spectrum into low- and high-frequency bands (Figure 2a, middle). Importantly, this simple description of amplitude coupling in terms of one or two states was very accurate (goodness-of-fit > 96% across datasets; Supplementary Results S3). The entire connectome of amplitude coupling thus evolves as a whole from rest to task.

**Figure 2:**
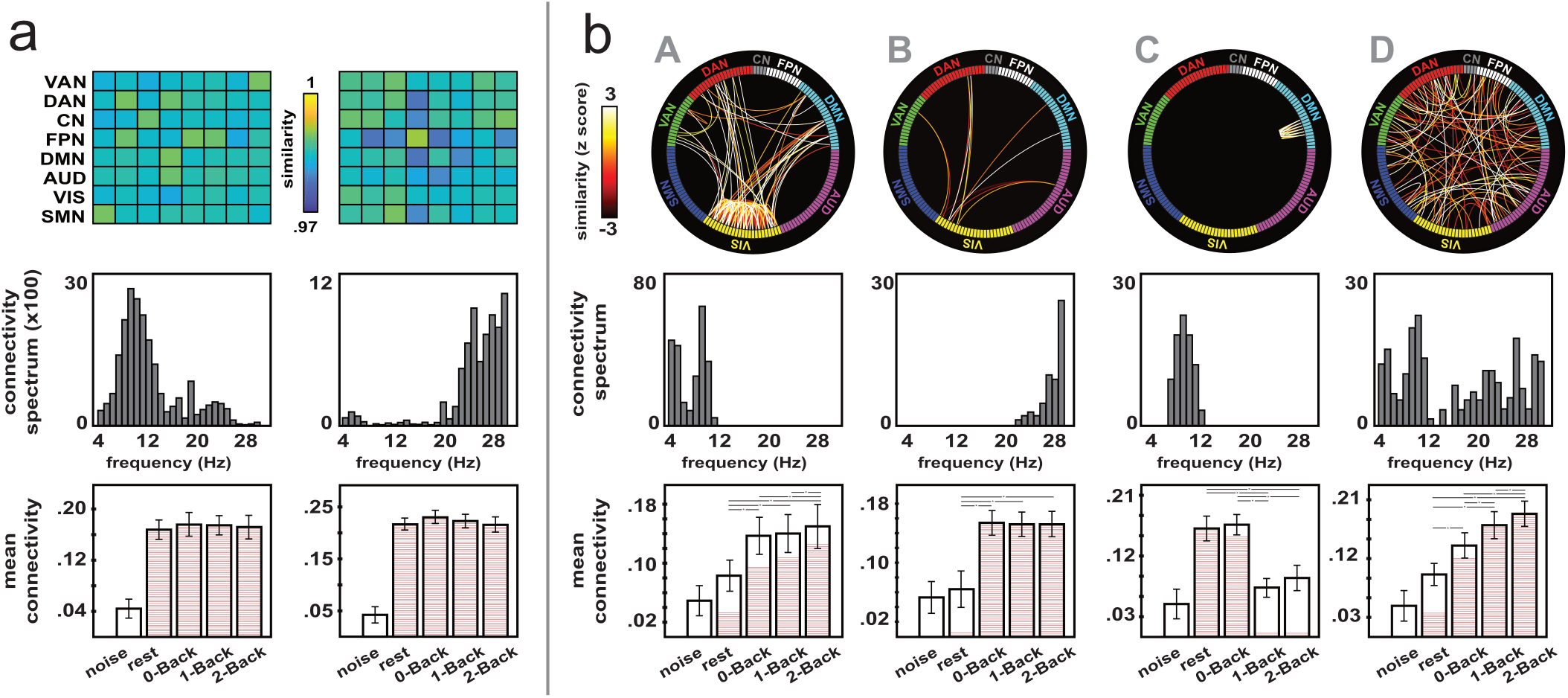
Connectivity states for the *n*-Back experiment. **a**, Amplitude correlation. **b**, Phase coupling. Each column corresponds to one state, for which we show different characteristics: **top**, connectivity map; **middle**, spectral distribution (number of connections in the state per frequency bin); **bottom**, mean state connectivity (i.e., average across all couplings in the state) per condition, noise estimate included. Links in these connectivity maps were weighted by a measure of similarity between their task-dependent functional connectivity pattern and the state pattern shown at the bottom. For visualization purposes, the amplitude correlation state maps were summarized using coarse network-level matrices due to the overwhelming number of links to draw. For phase coupling states, the similarity values were converted into *z* scores to better emphasize color contrasts. The state connectivity bar plots at the bottom show the group mean and SEM across single-subject values. The filling in each bar indicates the proportion of state links that are significantly above noise level in the corresponding condition (*p* < 0.05 corrected for the false positive rate). Stars indicate significant differences across two conditions (*t* tests, *p* < 0.05, noise condition excluded). VAN, ventral attentional network; DAN, dorsal attentional network; CN, control-executive network; FPN, fronto-parietal network; DMN, default-mode network; AUD, auditory network; VIS, visual network; SMN, sensorimotor network.

To characterize how amplitude coupling depends on task, we took advantage of the accuracy of state classification and assessed the task-related evolution of mean state connectivity. Figure 2a (bottom) shows the evolution of low- and high-frequency amplitude correlation from rest through the three *n*-Back conditions, and suggests that neither is modulated by task or perceptual/cognitive load. To test this hypothesis statistically for each state of each experiment, we estimated mean state connectivity at the individual level via dual regression of the clustering model and applied a one-way ANOVA with condition (e.g., rest, 0-Back, 1-Back, and 2-Back) as factor. The test was non-significant for all states (*F* < 3.8, *p* > 0.33 across states and datasets) with very low effect size (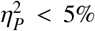 of variance explained by task-related changes; Figure 3, light grey). Given the high accuracy of state classification and the wide variety of experiments, we conclude that amplitude coupling is largely dominated by intrinsic integration.

**Figure 3:**
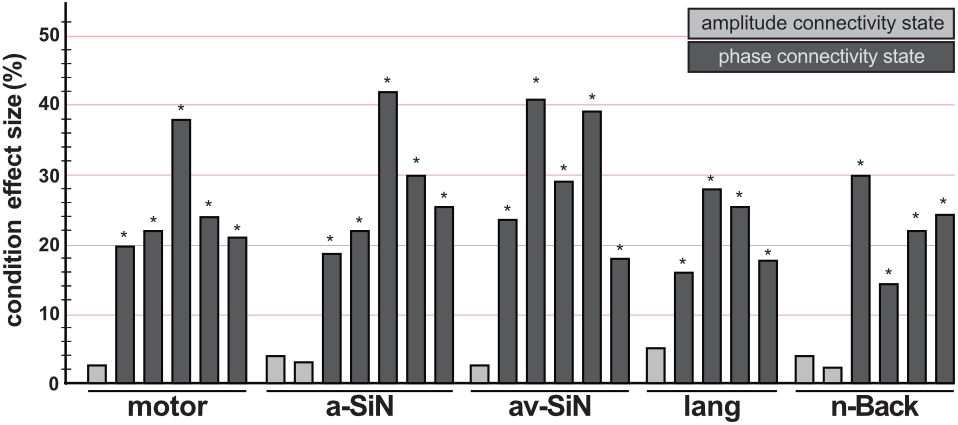
Task (in)dependence of connectivity states. The effect sizes of the ANOVA applied to state connectivity patterns (see Figure 2, bottom, for the *n*-Back states) are gathered across all our experiments. Each bar corresponds to one state (light grey, amplitude correlation; dark grey, phase coupling). The effect size measures the fraction of variance explained by task-related changes in group-level state connectivity. Stars indicate significance of the corresponding ANOVA at *p* < 0.05. motor, motor sequence learning; a-SiN, auditory speechin-noise comprehension; av-SiN, audiovisual speech-in-noise comprehension; lang, covert language production.

### Task-dependent states of phase coupling

Phase connectivity exhibited more diverse task dependence, with 4 to 5 states required for optimal classification (goodness-of-fit > 93% across datasets; Supplementary Results S3). Figure 2b details the states for the *n*-Back experiment. We identified a low-frequency (< 11 Hz; state A) and a high-frequency (> 23 Hz; state B) visuo-attentional state, both involving phase coupling within and across the visual, the ventral/dorsal attentional, and the default-mode (DMN) networks. The high-frequency state emerged during task only (not at rest) and was not modulated by working memory load, whereas the low-frequency state was partially active at rest but its phase coupling strength increased with working memory load (Figure 2b, bottom). We also observed an intra-DMN state in the alpha band (8 - 13 Hz; state C) that was recruited at rest and during the simple 0-Back condition but got disconnected during conditions involving higher working memory load (1- and 2-Back). One last, broadband state appeared spread across the connectome (state D) but disclosed steadily increasing phase coupling from rest to the 2-Back condition. Similar results held after eliminating task event-related responses (Supplementary Results S4, which includes further discussion).

Meaningful states of phase coupling also emerged in the other experiments (Supplementary Results S2, for details). Briefly, all phase connectivity states except one involved coupling increases from rest to task and from lower to higher perceptual/cognitive load, and they were maximally recruited during task. Most states consisted of connections between networks that were functionally specific to the task at hand (motor sequence learning, visual and motor; speech-in-noise comprehension and language production, auditory/language and motor) and attentional networks. The motor and speech-in-noise datasets also evidenced states involving the DMN and attentional networks with connectivity increasing from rest to task. Finally, one language production state disclosed resting-state intra-DMN phase synchronization that decreased from rest to task, similarly to the *n*-Back state C.

We confirmed statistically this qualitative description of task-dependent phase couplings using an ANOVA on mean state phase connectivity. In stark contrast with amplitude correlation, all states were significantly modulated by task (*F* > 11.3, *p* < 0.0011, 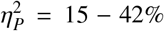 across states and datasets; Figure 3, dark grey), including those active at rest. Phase coupling thus appears extrinsic. The reproducibility of our results across all our five datasets, which encompassed distinct functional systems, shows that results did not depend on a specific cognitive task. They also proved resilient to methodological changes in connectivity estimation (spectral range, brain parcellation, power regression, leakage correction, elimination of event-related responses; Supplementary Results S4).

### Amplitude correlation modulates phase coupling

The fact that amplitude correlation is intrinsic prompts the question of its functional relevance. To answer this question, we focused again on the frequency-specific connections significantly exceeding phase-locking noise level and examined the inter-dependence of their resting-state amplitude coupling and their phase coupling at rest or during task. A substantial proportion of those connections disclosed significant, positive correlations (45 - 65% of Spearman correlation tests computed over subjects for each considered frequency-specific connection at *p* < 0.05 corrected for multiple comparisons; range across conditions and datasets). Accordingly, their distribution was heavily shifted towards positive correlations (Figure 4a; non-parametric test on distribution mean, *p* < 10^−7^). These correlation distributions were similar across tasks, suggesting that this partial inter-dependence of amplitude and phase coupling^11^ is intrinsic. For a more functionally relevant relationship, we also correlated resting-state amplitude coupling and phase connectivity *changes* from rest to task. Their distribution was now shifted towards negative correlations (Figure 4b), less prominently (9-14% of significant correlations) but still significantly for the mean (*p* < 10^−4^). Thus, connections with strong resting-state amplitude correlation tend to exhibit a higher baseline of phase connectivity and smaller phase coupling changes for task processing. We conclude that intrinsic integration as encoded in amplitude correlation facilitates the formation of extrinsic phase coupling.

**Figure 4:**
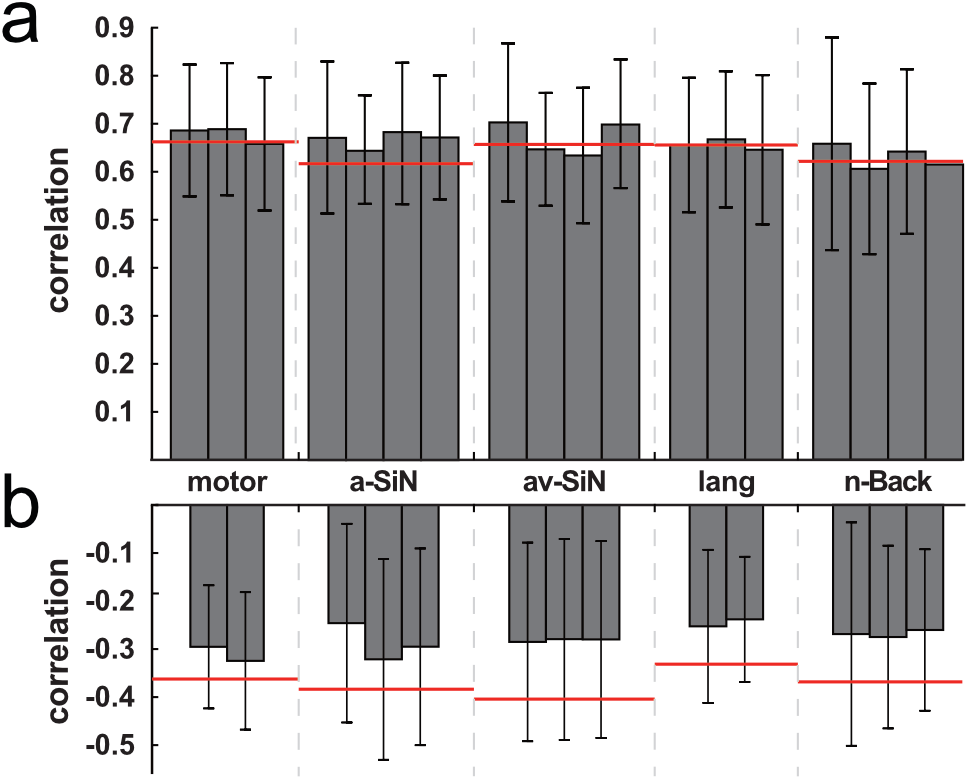
Correlation between amplitude and phase coupling. **a**, Mean and SD of Spearman correlations between resting-state amplitude correlation and phase coupling in each condition of our five experiments. Each bar corresponds to one condition. Only the connections that were included in phase connectivity state classification contributed to this distribution. Thresholds for correlation significance (*p* < 0.05 corrected for the false positive rate) are indicated in red. **b**, Similar plot for the correlation between resting-state amplitude correlation and phase coupling changes from rest to condition. motor, motor sequence learning; a-SiN, auditory speech-in-noise comprehension; av-SiN, audiovisual speech-in-noise comprehension; lang, covert language production.

### Phase coupling exhibits intrinsic dynamics

This facilitation effect suggests that amplitude coupling might induce intrinsic features in phase coupling. We explored this idea using the temporal development of amplitude correlation and phase locking (estimated over 10 s-long sliding windows^23^). Short-time amplitude and phase coupling fluctuations were positively correlated in time (Supplementary Results S5, which also identifies their temporal standard deviation as intrinsic). This extends the facilitation effect to finer timescales. We then relied on this temporal regression to isolate the part of time-averaged phase connectivity explained by amplitude coupling dynamics. This amplitude-based model of phase coupling was organized into RSNs (Figure 5a and Supplementary Results S1) and state classification revealed a single, task-independent state (goodness-of-fit > 94% across datasets, Supplementary Results S3; *F* < 2.1, *p* > 0.35, 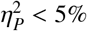 across conditions and datasets, Figures 5b,c). This does not contradict our earlier results since this contribution to time-averaged phase connectivity was subdominant (explained variance < 11%). Besides its task-related modulations, extrinsic phase coupling thus also exhibits intrinsic modulations closely tied to amplitude correlation dynamics.

**Figure 5:**
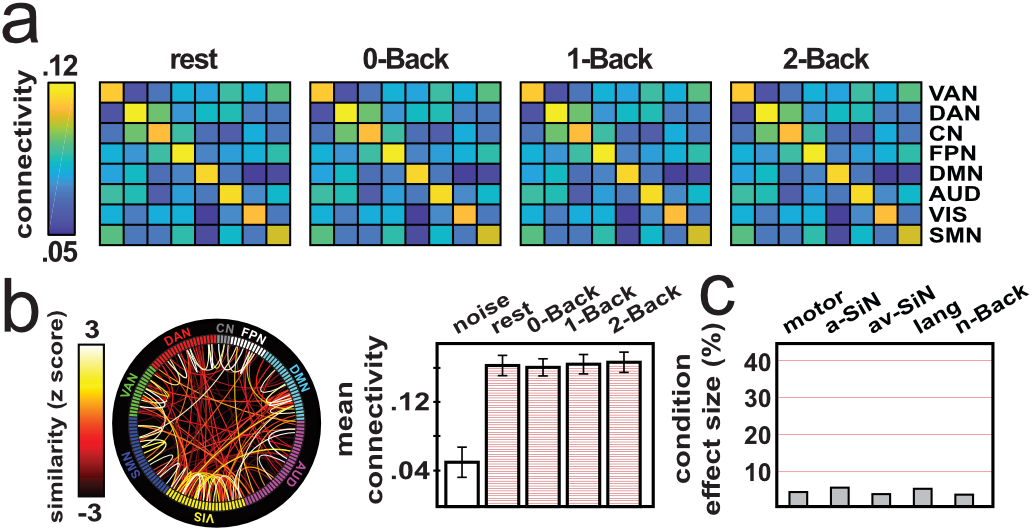
Amplitude-based model of phase connectivity. Analysis of the part of time-averaged phase coupling explained by amplitude correlation dynamics. **a**, Illustration of the broadband network-level connectivity across the *n*-Back conditions. **b**, Connectivity state obtained in the *n*-Back experiment. **c**, Effect sizes of the ANOVA applied to state phase connectivity across all experiments. VAN, ventral attentional network; DAN, dorsal attentional network; CN, control-executive network; FPN, fronto-parietal network; DMN, defaultmode network; AUD, auditory network; VIS, visual network; SMN, sensorimotor network; motor, motor sequence learning; a-SiN, auditory speechin-noise comprehension; av-SiN, audiovisual speech-in-noise comprehension; lang, covert language production.

## Discussion

Our unique MEG dataset comprising 141 adult subjects and 14 experimental conditions provides invaluable insights into the conception of functional brain organization^5^. The importance of intrinsic activity was recognized rapidly after discovering the DMN with positron emission tomography (PET) and then other RSNs with fMRI.^2,3^ Its study has since been dominated by a dichotomy between task-related and resting-state neuroimaging. However, the resting state is not task free but an introspective (albeit ill-controlled) condition alternating conscious processing of inner and external stimuli with spontaneous cognition (e.g., mind wandering).^7^ We advocate here a novel conceptual paradigm wherein intrinsic and extrinsic brain communications work in parallel and are dissociated by electrophysiological couplings rather than experimental conditions. Previous works emphasized the coexistence of intrinsic and extrinsic brain systems^25^ but without fully separating or characterizing them. We demonstrated that amplitude correlation is predominantly intrinsic, whereas phase coupling is extrinsic but still endowed with intrinsic dynamics closely tied to that of amplitude connectivity. The distinctive temporal resolution of MEG^18^ was essential to unveil this dissociation to which PET and fMRI are blind.^26^ This duality prompts the question of how they can coexist on a common anatomical connectivity substrate while showing strikingly different functionalities. This can be answered by considering plausible underlying neural mechanisms. The role of phase connectivity in network-level neural processes is understood in terms of neural firing synchrony.^16,17^ Phase locking gauges the tendency that pre-synaptic action potentials from one neural population arrives at the time of maximal post-synaptic potential in the other, thus favoring synchronous firing and information transfer.^17^ With hindsight, its extrinsic character appears unsurprising since firing patterns largely govern neural computations^27^. The mechanisms for amplitude coupling are less understood. Amplitude correlation presumably measures the tendency of neural populations to simultaneously exhibit high levels of excitability.^6^ Fluctuations in neural excitability are thought to be related to slow, endogenous variations in biophysical properties of neurons,^28^ e.g., extracellular potassium concentration^29^. They do not affect neural firing directly, which explains the intrinsic character of amplitude correlation, but indirectly modulate it.^28^ Cooccurrence of high excitability favors firing in one population upon incoming action potentials from the other and thus reinforces their phase synchronization. This phenomenon of phase-coupling facilitation through amplitude correlation provides a neurophysiological rationale for our identification of an intrinsic component of phase connectivity and its close interplay with amplitude connectivity dynamics^30^. In fact, it suggests that amplitude coupling (i.e., excitability covariation) drives intrinsic phase coupling through modulatory effects. It further supports recent work on highly transient phase coupling at rest^15^, although amplitude and phase connectivity were entangled there. Our approach was specifically designed to disambiguate them, at the expense of lower temporal resolution. Conversely, sustained phase synchronization affects neural excitability through, e.g., spike-timing dependent plasticity, and may thus slowly reorganize amplitude correlations.^6^ This suggests that amplitude coupling is intrinsic only so long as plastic changes are negligible. This was presumably the case in our experiments spanning 10 - 20 minutes. On the timescale of hours and longer, experience- and learning-dependent plasticity builds up^31^ and may reorganize intrinsic couplings.^3^ For example, memory con-solidation appears to modulate amplitude connectivity after motor sequence learning.^32^ Still, short-term plasticity in visual cortices^33^ might explain minute-scale topological changes in amplitude correlation during, e.g., movie watching^34^.

This neurophysiological perspective triggers a demotion of resting-state studies to characterize intrinsic activity^35^. The fact that intrinsic integration is organized into RSNs explains why they are observable at rest, but it is not limited to RSNs since amplitude correlation covered the whole connectome. More strikingly, resting-state phase connectivity revealed an extrinsic RSN, i.e., the phase-synchronized DMN, which further supports the idea that resting-state connectivity is not exclusively intrinsic. Phase coupling among DMN regions was previously postulated from indirect considerations about DMN mapping with MEG amplitudes.^36^ Here, we go further and claim (alongside recent fMRI data^37^) the existence of two functionally distinct modes of DMN integration. In contrast to the intrinsic DMN, the extrinsic DMN exhibited decoupling during tasks with high cognitive load (language production, 1- and 2-Back). This is akin to the goal-directed PET/fMRI deactivations that historically defined the DMN.^38^ This analogy actually carries further. Phase DMN desynchronization was accompanied by increased synchrony in attentional networks, which is reminiscent of the fMRI anticorrelation between task-positive/task-negative networks^39^. Additionally, some DMN nodes increase their PET/fMRI activity in tasks such as speech comprehension,^40^ during which DMN phase connectivity strengthened. These findings suggest that extrinsic phase coupling represents a better neural correlate of task-evoked PET/fMRI modulations than the slow amplitude fluctuations usually considered in this context^26,41^. Another, more practical consequence is that the classical division between resting-state and task-related neuroimaging should be alleviated and replaced by an informed choice of the relevant neurophysiological coupling type. Phase connectivity should be used to study active neural networks during stimulation or task performance, but also at rest to characterize fleeting spontaneous brain processes, which are currently little understood^7^. The resting state is generally used to compare physiological or pathological conditions and, critically, assumes the absence of task performance biases.^42^ Extrinsic phase coupling should undergo such a bias, even at rest, as different subtypes of spontaneous cognition transiently reconfigure RSNs^7^. Further, physiological processes such as ageing and brain disorders do affect spontaneous cognition.^43–46^ In this light, findings of age- or disease-related changes in phase connectivity^47,48^ may need careful reconsideration. Amplitude correlation appears better suited to avoid these biases and identify reorganizations of intrinsic functional integration. Notwithstanding, phase connectivity may still be used to reveal alterations in spontaneous brain processes. We thus suggest that studies systematically disentangle intrinsic and extrinsic aspects of resting-state activity when comparing physiological processes or brain disorders.

In conclusion, the intrinsic/extrinsic duality of functional integration revealed in our study provides a novel conceptual framework to human brain organization. This framework is firmly grounded in the electrophysiology of long-range neural communication. Ultimately, it may prove invaluable to cognitive and clinical applications of brain network science.

## Methods

Experimental and analytical procedures along with associated references are available in the Supplementary Information.

## Acknowledgements

This study was supported by the Fonds de la Recherche Scientifique (FRS-FNRS, Brussels, Belgium; PDR 3.4558.12, EOS “MEMODYN”), the Wiener-Anspach Foundation (Brussels, Belgium) which also funded M.S., the Fonds Erasme (Brussels, Belgium; research conventions “Les Voies du Savoir” and “Marc Errens”) which also funded M.S. (convention “Marc Errens”), A.D., M.N., and M.V.G. The program Attract of Innoviris supported M.B., F.D., and J.B. (grant 2015-BB2B-10). M.B. also acknowledges funding from the Spanish Ministry of Economy and Competitiveness (Spain, grant PSI2016-77175-P) and the Marie Skłodowska-Curie Action of the European Commission (grant 743562). The Fonds Wetenschappelijk Onderzoek (FWO, Brussels, Belgium) supported L.C. (aspirant grant 11B7218N), J.V.S. (“Krediet aan Navorser”, 1501218N), and G.N. (“Fundamenteel klinisch mandaat”, 1805620N). T.C. is Clinical Master Specialist Applicant to a PhD and X.D.T. is Postdoctorate Clinical Master Specialist at the FRS-FNRS. J.B. is supported by the Fonds Marguerite-Marie Delacroix. G.N. acknowledges grants from the Belgian Charcot foundation and Genzyme-Sanofi. M.W.W. is supported by the NIHR Oxford Health Biomedical Research Centre, the Wellcome Trust (106183/Z/14/Z, 203139/Z/16/Z, 215573/Z/19/Z), and the New Therapeutics in Alzheimer’s Diseases (NTAD) study (funded by UK MRC and the Dementia Platform UK). The MEG project at the CUB Hôpital Erasme is financially supported by the Fonds Erasme (convention “Les Voies du Savoir”).

## Author contributions

M.S.: study design, data collection, methodological developments, data analysis, data interpretation, writing paper. M.B., L.C., A.D., T.C., L.R., F.D., J.B., M.N., M.V.G, J.V.S.: data collection, writing paper. G.N., C.U., P.P., S.G., M.W.W.: data interpretation, writing paper. X.D.T.: conceptualization, study design, data analysis, data interpretation, writing paper. V.W.: conceptualization, study design, methodological developments, data analysis, data interpretation, writing paper.

## Supplementary Information

### Methods

#### Subjects and experimental procedures

To demonstrate the generality of our analysis, we considered five experiments targeting different types of brain systems (e.g., motor, auditory, language, or working memory). Each dataset was acquired in a different group of healthy adult subjects. All were right-handed (Edinburgh handedness inventory test^1^) with no history of neurological or psychiatric disease, and no auditory deficit for the speech-in-noise comprehension experiments. They participated after signing a written informed consent. See Table S1 for demographic information.

**Table S1:** Demographic information. *N*, number of subjects; motor, motor sequence learning; a-SiN, auditory speech-in-noise comprehension; av-SiN, audiovisual speech-in-noise comprehension; lang, covert language production.

| | *N* | age (years, mean ± SD) | sex (female:male) |
| --- | --- | --- | --- |
| motor | 26 | 23.2 ± 2.7 | 9:17 |
| a-SiN | 25 | 30.1 ± 3.2 | 12:13 |
| av-SiN | 25 | 23.0 ± 2.9 | 15:10 |
| lang | 30 | 32.5 ± 9.1 | 16:14 |
| *n*-Back | 35 | 43.2 ± 9.2 | 22:13 |

Each experiment included a resting state as well as several attentional and cognitive tasks in which perceptual and cognitive loads varied in each functional modality, as further detailed below. During the resting-state conditions, subjects were instructed to relax with their gaze fixated either at a point on the wall of the magnetic shielded room (where the MEG recordings took place) or at a cross on a MEG-compatible screen. The resting state was systematically obtained as a session separate from the other tasks. The different tasks of an experimental paradigm were either performed sequentially as different sessions (motor sequence learning and auditory speech-in-noise comprehension) or intermingled within one uninterrupted session using a block design (audiovisual speech-in-noise comprehension, language production, and *n*-Back). Each task lasted about 5 minutes in total. The ordering of the rest and task conditions varied across the experiments. Subjects were seated on the MEG armchair, except in one experiment (audiovisual speech-in-noise comprehension) in which subjects lied on a MEG-compatible bed. The convergence of our main results indicates that they were not affected by these differences in experimental procedures or MEG data collection. To ensure the reproducibility of our study, we describe below these procedures for each of the five experiments reported in the main text. We had prior approval by the National MS Center Melsbroek and UZ Brussel ethics committees for the *n*-Back experiment, and by the CUB Hôpital Erasme ethics committee for the others.

##### (i) Motor sequence learning

This paradigm targeted the motor system and the learning of a new finger motor sequence (i.e., procedural learning). The experiment contained a restingstate session (cross fixation) followed by a simple left finger tapping task and then by a left finger sequence learning task. These tasks were performed in two separate sessions and consisted in the reproduction of a pattern of key presses on a MEG-compatible four-button box (fORP, Current Designs Inc.). The simple finger tapping task required simultaneously pressing all buttons with four fingers (index to little finger) every 5 seconds upon an auditory cue. In the sequence learning task, subjects had to reproduce a complex sequential pattern (4-1-3-2-4, where 1 corresponds to the index and 4 to the little finger) as fast and accurately as possible. These procedures were adapted from a previous MEG experiment used to study motor sequence learning.^2–4^

##### (ii) Auditory speech-in-noise comprehension

This experiment targeted the brain systems involved in connected speech processing in the presence of informational noise^5^ with varying intensity, which modulates subjects’ ability to understand speech. It consisted in one resting-state (wall fixation) and three listening conditions performed in separate, randomly ordered sessions. To each listening condition was assigned a noise level (signal-to-noise ratio: noiseless, −5, and 5 dB) and a story that subjects had to attend to. The stories were randomly selected from a set of six recordings recounted by different French-speaking readers (sex ratio 3:3) and obtained from an audiobook database (http://www.litteratureaudio.com). Informational noise was built as a multi-talker background of six French speakers (sex ratio 3:3) talking simultaneously, and added to these recordings at a specified signal-to-noise ratio. This audio material was delivered at an average intensity of 60 dB through a MEG-compatible 60 × 60 cm^2^ flat-panel loudspeaker (SSH sound shower, Panphonics) placed in front of the subjects. Subjects were instructed to fixate a wall point and to try and attend the story. Further background on this paradigm and details on its use to identify slow-wave cortical oscillations tracking the syllabic and phrasal speech content, are described in previous publications.^6,7^

##### (iii) Audiovisual speech-in-noise comprehension

This experiment was similar to the previous one, except that it focused on the impact of noise type (i.e., non-informational vs. informational^5^) rather than noise intensity, and included visual support (speaker’s lips reading) to facilitate speech-in-noise comprehension^8^. Here, four sessions of the audiovisual task preceded the resting state (fixation cross). The audiovisual material was randomly selected from twelve different recordings of four French-speaking actors telling a story. Each session corresponded to one story told by a different actor, with various types of speech noise occurring in randomly-ordered, 30-second blocks; of which two were noiseless, two with non-informational noise, and two with informational noise. Non-informational noise consisted in a Gaussian white-noise signal filtered between 100 and 10^4^ Hz for one of the corresponding blocks, and in a similar signal whose spectrum was modulated to match that of the speaker’s voice in the other block. Informational noise consisted in a multi-talker background of five French speakers of same gender than the speaker in one block, and of opposite gender in the other. Informational noises interfere with speech comprehension significantly more than non-informational noises.^5^ Audio and video materials were presented time locked using a screen seen by the lying subjects through a mirror placed above their head, and a 60 × 60 cm^2^ flat-panel loudspeaker (SSH sound shower, Panphonics; 60 dB intensity), both MEG-compatible and placed in front of the bed. An extended version of this paradigm was used to investigate the relationship between slow-wave brain tracking of speech, noise type, and reading abilities in children and dyslexia.^9^

##### (iv) Covert language production

This experiment targeted the functional systems associated with verbal language production. It consisted in a resting-state session (fixation cross) followed by two covert verbal language production tasks executed within a single session in a block design: a picture naming task where subjects had to silently name an image presented on screen, and a verb generation task where they had to silently generate verbs associated to such an image. The latter entails a higher cognitive load than the former.^10^ For each subject, 40 images were randomly selected among 80 images taken from two databases, one dedicated to picture naming^11^ and the other to verb generation^12^. The two tasks were presented in a total of 20 alternating 30-second blocks. Each block was preceded by a one second-long cue identifying the task to perform (the choice for the initial block being random) followed by two seconds of cross fixation before block start. Each block consisted in the presentation of eight images shown during one second on a MEG-compatible screen, with random inter-stimulus intervals (cross fixation during 2.7 ± 0.8 s, mean SD) constrained to sum to 22 seconds within the block.

##### (v) Multi-item visual *n*-Back

This classical paradigm targeting visuo-attentional processes and working memory was performed at three levels of difficulty (*n* = 0, 1, 2) to modulate attention and working memory load.^13^ This experiment contained a resting-state session (wall fixation) and one continuous *n*-Back session in which the three levels were presented by block. Each level was assigned to four blocks, for a total of 12 blocks, with random ordering except that the same level could not occur twice consecutively. Each block started with task instructions shown during 15 seconds on a MEG-compatible screen, followed by the presentation of 20 visual items (here, letters) projected on the screen for one second and separated by an inter-trial interval of 2.8 seconds. Subjects were instructed to press a key on a MEG-compatible button box when the current item corresponds to the letter ‘X’ (*n* = 0), the previous item (*n* = 1), or the one before (*n* = 2). This setup has been used in conjunction with MEG to characterize extensively the brain electrical evoked and rhythmical activity involved during this visual *n*-Back.^14^

#### Data acquisition

Electrophysiological brain activity generates small extracranial magnetic fields (of the order of 10^−15^T) that are measurable using MEG based on superconducting quantum interference devices (SQUIDs).^15,16^ All MEG recordings in this study were obtained with a 306-channel, whole-scalp-covering SQUID neuromagnetometer (Neuromag Vectorview™/Triux™, MEGIN, Helsinki, Finland) placed in a light-weight magnetically shielded room with active internal compensation of remnant slow magnetic drifts (Maxshield™, MEGIN)^17^. They were collected using either a Vectorview system (auditory speech-in-noise comprehension), a Triux system (motor sequence learning, audiovisual speech-in-noise comprehension, and covert language production), or both (*n*-Back; Vectorview: 10 subjects, Triux: 25 subjects) due to an upgrade from Vectorview to Triux. These two neuromagnetometers have the same sensor array, organized into 102 triplets of one magnetometer and two orthogonal planar gradiometers,^15,16^ with some differences in sensor dynamic range and magnetic environment. Still, their noise levels were comparable, especially after environmental noise cancellation. Previous works that mixed Vectorview and Triux recordings did not reveal any significant change in data quality for task-related activity^18,19^ or resting-state functional connectivity^20,21^. The MEG acquisition electronics comprised an analog band-pass filter (0.1-330 Hz) and digital conversion at the sampling rate of 1 kHz. Head movements within the MEG helmet (which is not attached to the subjects’ head) were tracked using four position indicator coils continuously emitting a localizable high-frequency (around 300 Hz) magnetic field during the recordings. For the paradigms involving a block design (audiovisual speech-in-noise comprehension, covert language production, and *n*-Back), trigger signals were also sent to the MEG acquisition electronics to identify the start, the type of condition, and the ending of each block.

A standard 3D T1-weighted cerebral magnetic resonance image (MRI) of each subject was further acquired after the MEG sessions, using a 1.5 T Intera™ scanner (Philips, Best, The Netherlands) for the motor sequence learning and auditory speech-in-noise comprehension experiments, a hybrid 3 T Signa™ PET-MR scanner (General Electrics Healthcare, Milwaukee, Wisconsin, USA) for the audiovisual speech-in-noise comprehension and covert language production experiments, and a 3 T Achieva™ scanner (Philips) for the *n*-Back experiment. The location of fiducials, position indicator coils, and over 300 head-shape points were digitized to enable the coregistration of MEG and MRI coordinate frames.

#### Data preprocessing and spectral decomposition

From thereon, we processed in a similar way all continuous MEG data recorded during the different sessions (i.e., before their separation into blocks). Signal space separation^22^ was used to suppress environmental magnetic interferences and correct for head motion on the basis of the indicator position coils (Maxfilter™ v2.2, MEGIN). We then eliminated physiological magnetic interferences induced by cardiac and ocular activity as well as electronic artefacts, with an independent component analysis (FastICA algorithm with pre-whitening, dimension reduction to 30, and nonlinearity contrast function tanh)^23^ of the MEG data filtered beforehand between 0.5 and 45 Hz. The components corresponding to these artefacts were identified and regressed out of the full-rank data^24^ (number of removed components: 3.8 ± 1.2, mean ± SD across sessions, subjects, and experiments; range: 3 - 6). The cleaned continuous MEG signals were then decomposed spectrally by filtering them in narrow, 1 Hz-wide non-overlapping frequency bins covering the 3.5 - 30.5 Hz interval. We chose this interval because it covers the theta (4-8 Hz), alpha (8-12 Hz) and beta (12-30 Hz) bands that support the spectral content of MEG RSNs^25^, and because the MEG signal is much noisier in delta (< 4 Hz) and gamma (> 30 Hz) frequencies^16^. The analytic MEG signals were extracted with Hilbert transformation to obtain a time-frequency decomposition amenable to frequency-specific amplitude and phase connectivity analysis.

#### Reconstruction of brain electrical activity

Neuromagnetic activity as measured with MEG is a blurred version of the underlying neural electrical currents, because magnetic fields spread in the course of their propagation from the brain to extracranial sensors. For this reason, reconstructing these currents from MEG data is an ill-posed problem that requires a regularized inversion of this field spread.^15^ The neuromagnetic field spread was estimated using MEG forward modeling, i.e., a numerical computation of the magnetic field generated by any current source distribution and propagating through head tissues. We computed each forward model individually using the single-layer boundary element method (MNE-C suite^26^). This approach provides an accurate approximation of the neuromagnetic field spread by considering head conductivity as being homogeneous inside the brain volume and vanishing outside.^15^ Individual, realistically-shaped brain volumes were obtained from tissue segmentation of their anatomical MRI (Freesurfer^27^), and their geometric position with respect to MEG sensors were determined by manual MEG-MRI co-registration (MRIlab™, MEGIN). The forward model was computed for three orthogonal current sources placed at each node of a grid sampling the brain volume. This grid was built from a cubic, 5-mm grid cropped within the Montreal Neurological Institute (MNI) template MRI brain volume and normalized onto each subject’s MRI (SPM12)^28,29^. This allows to directly work with MNI nodes and facilitates group analysis. In particular, the nodes corresponding to the RSN-based brain parcels used in this work (Figure 1a; see Figure S1 for more detailed views) could be directly identified from their MNI coordinates.

The current source distribution of brain oscillatory activity was then reconstructed in the whole brain volume grid by minimum norm estimation, i.e., a regularized inversion of the MEG forward model.^15,30^ This choice of reconstruction algorithm was based on a recent work demonstrating that full DMN imaging with MEG amplitude correlation requires minimum norm estimation instead of the popular beamforming approach.^31^ The regularization was based here on a frequency-specific noise covariance estimated from empty-room MEG recordings (5 minutes, with environmental noise cancellation and spectral decomposition similar to the resting-state/task data), and a regularization parameter adapted to the frequency-specific MEG signal-to-noise ratio via the prior consistency condition^32^. The reconstructed current sources were finally projected onto their direction of maximum variance^33,34^ so as to fix optimally their orientation and enable the usage of bivariate connectivity measures. Of note, in the context of amplitude correlation analysis, an alternative sometimes used is to work with the Euclidean norm of three-dimensional current sources.^35,36^ Both approaches yield very similar connectivity estimates.^31^ Source power was computed as the signal variance of these projected currents with proper noise standardization^37^ to account for the depth bias.

#### Frequency-specific functional connectivity

We estimated functional connectivity between the reconstructed MEG oscillatory activity at each pair of nodes in the brain parcellation (Figures 1a and S1) and in each frequency bin. Connectivity measures are afflicted by the spatial leakage effect, i.e., spurious coupling inflation due to the typical spatial smoothness of minimum norm estimates (which is eventually rooted in magnetic field spread).^38^ We controlled for spatial leakage using a geometric correction that models the leakage effect at one (target) node from another (seed) node based on the MEG forward model, and then subtracts it from the target before connectivity estimation.^32^ Compared to other, popular corrections based on signal orthogonalization,^25,39^ this method offers the advantage of preserving zero phase-lag synchronization and also suppresses the leakage screening effect (i.e., a spurious coupling reduction) recently reported^40^. This preservation was *a priori* deemed important for our study, especially given the expectation of zero-lag linear synchronization within the DMN^31^. (See cross-correlation of amplitude and phase connectivity, for an additional benefit of the geometric correction.) Amplitude coupling was computed as the temporal Pearson correlation between the Hilbert envelopes of the seed and leakage-corrected target signals.^25,33,34^ Phase coupling was computed using the phase-locking value^41^ based on the instantaneous phase difference of their analytical signals.^42^ For each subject and experimental condition, time-averaged connectivity (used in the bulk of our analyses) was estimated over the entire time periods corresponding to the condition. Their temporal development was extracted by measuring connectivity in short, 10 s-long windows sliding across these time periods by steps of 5 s.^21,43^ For both coupling types (and each time window when appropriate), connectivity estimates across all seed-target pairs were gathered in a node-by-node connectivity matrix, which was further averaged with its transpose to account for slight asymmetries induced by leakage correction^25^. The alternative would be to use the symmetrical multivariate orthogonalization^44^ that avoids this issue. However, it was inapplicable in our case since it is limited to sparser brain parcellations. Finally, given that power changes may reflect modifications in signal-to-noise ratio and thus bias connectivity estimation,^45^ each node-to-node connectivity measure was further corrected by regressing out source power at the two corresponding nodes.

The significance of time-averaged connectivity measures was assessed statistically by one-sided paired *t* tests against their noise level (estimated from empty-room MEG recordings). Given the multiple comparisons involved (11935 links and 30 frequency bins to test) and that each experiment contained several conditions (one resting state and 2 - 3 tasks), we relied on a maximum statistic^46,47^ (here, the maximum *t* value taken over all connections, frequencies, and conditions) to detect frequency-specific couplings significantly above noise level in at least one experimental condition while controlling the false positive rate at *p* < 0.05. To perform the test, we generated non-parametrically 2000 null samples of this statistic under the hypothesis that all conditions only disclosed connectivity noise. Each sample was simulated by exchanging, independently in each condition and for a random selection of subjects, the connectivity data obtained in the said condition with the corresponding noise estimates. We then located significant couplings by masking out all connections disclosing a univariate *t* value below the 95^th^ percentile of these null samples in all conditions (resting state and tasks).^46,47^ We restricted all subsequent analyses to couplings within this mask.

**Figure S1:**
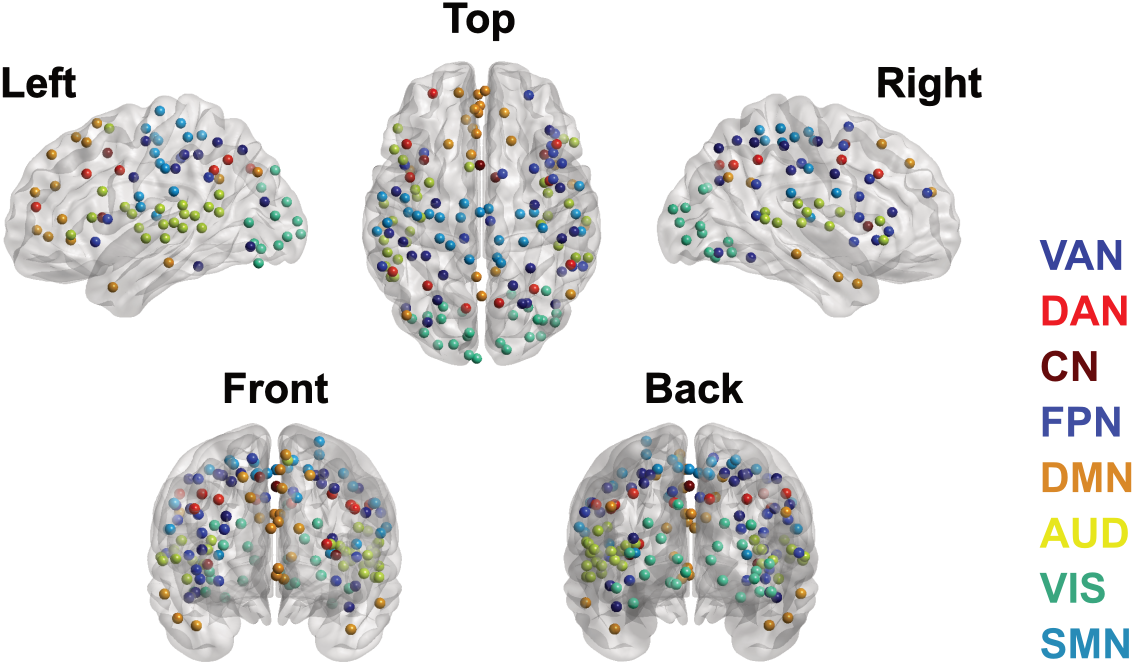
Network-based brain parcellation. Detailed location of the parcel nodes in the MNI brain, shown per RSN. We refer to Table 1 in Della Penna et al.^40^ for the corresponding list of MNI coordinates and labels. VAN, ventral attentional network; DAN, dorsal attentional network; CN, control-executive network; FPN, fronto-parietal network; DMN, default-mode network; AUD, auditory network; VIS, visual network; SMN, sensorimotor network.

#### Task-dependent connectivity state classification

We used Lloyd’s algorithm for *k*-means clustering^48^ to partition group-mean functional connectivity data across experimental conditions into distinct states. The distance measure between two task-dependent connectivity patterns, treated in the algorithm as *c*-vectors (where *c* denotes the number of experimental conditions), was taken as one minus the cosine of their angle. Accordingly, the angle cosine allowed to measure the degree of similarity between the task-dependent pattern of one coupling in a state and its mean state pattern. In our setup, each state corresponds to a subset of frequency-specific couplings. Task-dependent mean state connectivity was defined as group-mean functional connectivity averaged over this subset separately in each experimental condition. Individual mean state connectivity was inferred via dual regression, i.e., by performing this subset average independently for each subject. This data was subjected to a repeated-measure ANOVA to detect a statistically significant effect of the experimental condition on group-level mean state connectivity. The associated effect size was measured using Cohen’s partial eta squared 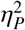.^49^

Building on the fact that clustering is a particular case of linear modeling, we assessed the goodness-of-fit of these state models as the fraction of explained variance in the connectivity data. Specifically, we built connectivity state models at the single-subject level by assigning the individual mean state connectivity value to each frequency-specific coupling in the corresponding state. Goodness-of-fit was then measured as the fraction of connectivity variance across subjects and conditions explained by this model data. One issue in state classification is that the number *k* of states to classify must be selected beforehand. This parameter is often chosen qualitatively by running state classification for a range of *k* values and drawing a so-called elbow curve to identify a compromise between high model goodness-of-fit (ensured at large *k*) and low model complexity (small *k*). See Results S3 for an illustration. Instead, we used here a robust method to select the optimal *k* value based on the gap statistics^50^. This approach also has the advantage of being able to detect situations where connectivity data are not separable into distinct states (*k* = 1), which was ideally expected for intrinsic integration (i.e., a single, task-independent state).

#### Cross-correlation of amplitude and phase connectivity

We examined the inter-dependence between time-averaged restingstate amplitude correlation and either (i) phase coupling at rest or in a task condition, or (ii) the difference between phase coupling during a task condition and phase coupling at rest. We used Spearman rank correlation computed across subjects and independently for each frequency-specific coupling. Analogously to the *t* tests described above (see frequency-specific functional connectivity), we controlled the false positive rate at *p* < 0.05 using a maximum statistic^46,47^, i.e., the maximum (case i) or the minimum (case ii) in the distribution of correlation values taken over all frequency-specific couplings and over all conditions (case i) or condition differences (case ii). Null samples were simulated by randomly reshuffling subjects ordering in the resting-state amplitude correlation data before re-computing this distribution (2000 permutations). We also tested whether the mean of this distribution was positive (case i) or negative (case ii) by extracting null samples of this mean statistic in the course of these simulations.

Crucially, our usage of the geometric spatial leakage correction^32^ when estimating functional connectivity was key to enable these correlation analyses. Signal orthogonalization mixes phase and amplitude signals^38^ and spuriously hides any inter-dependence between amplitude and phase couplings.^51^ This bias is not shared by the geometric correction.^38^

#### Amplitude-based model of phase coupling

We isolated the part of phase connectivity explained by amplitude correlation at the intra-subject level, by designing a linear regression model with sliding-window phase locking as dependent variable and sliding-window amplitude correlation as independent variable. This was done separately for each entry of the frequency-specific connectome, each subject, and each experimental condition. This temporal regression captured the dynamical relationship between phase and amplitude coupling. Usage of this model at the level of time-averaged connectivity allowed to single out the dynamic contribution of amplitude correlation to phase locking.

### Results

#### S1. Network-level functional connectivity plots (Figure S2)

In the main text, we used the *n*-Back data to illustrate the fact that time-averaged amplitude correlation (Figure 1c) and amplitude-based phase coupling (Figure 5a) are prominently structured in terms of RSNs, whereas time-averaged phase coupling is not (Figure 1d). Figure S2 shows similar data for the other four experiments considered in this study (motor sequence learning, auditory and audiovisual speech-in-noise comprehension, and covert language production) and demonstrates that these claims are reproducible.

#### S2. Further details on task-dependent connectivity states (Figures S3–S7)

Task-related connectivity state classification was illustrated in the main text with the *n*-Back dataset. We describe here the states that emerged from the motor sequence learning (Figure S3), auditory (Figure S4) and audiovisual (Figure S5) speech-in-noise comprehension, and language production (Figure S6) tasks. Throughout all datasets and consistently with the *n*-Back case (Figure 2a), all amplitude correlation states exhibited spatial homogeneity and task independence (as was shown in Figure 3). All phase connectivity states were task dependent (as again shown in Figure 3) but their spatial distribution was specific to the task at hand. Their maps are gathered in Figure S7, where we show them with node labels. All these states involved phase coupling increases from rest to task, except for the two intra-DMN states (i.e., *n*-Back state C, see Figure 2b, and language production state D, see Figure S6b).

##### (i) Motor sequence learning (Figure S3)

We identified two visuo-attentional states, one characterized by alpha-band phase coupling unmodulated across the two motor tasks (state A), and the other by low frequency (< 12 Hz) phase coupling increase from simple finger tapping to sequence learning (state B). The latter was reminiscent of the *n*-Back state A (Figure 2b). We also identified two motor-attentional states, with a similar discrimination in their task dependence (states C and D). State E involved most prominently the DMN and attentional networks and was characterized by a boost in phase coupling during sequence learning.

##### (ii) Auditory speech-in-noise comprehension (Figure S4)

This experiment revealed an auditory/language-motor-attentional state exhibiting higher theta-band (4 - 8 Hz) phase coupling in the presence of speech noise, but independently of noise level (state A). We also observed three states involving the DMN and attentional networks (states B–D), somewhat akin to the motor sequence learning state E. Their phase coupling was not clearly modulated by speech noise or its level. State E was less obvious to interpret.

##### (iii) Audiovisual speech-in-noise comprehension (Figure S5)

As in the purely auditory paradigm, we identified auditory/language-motor-attentional states (states A–C) and DMN/attentional states (states D and E). States A, C, and D were characterized by a phase coupling increase in the presence of speech noise, but independently of noise type. States B and E disclosed steadily increasing phase coupling and in particular stronger synchronization in the informational noise condition, which impedes speech comprehension more than then non-informational noise condition. These two noise type-dependent states were spectrally concentrated on the alpha band.

##### (iv) Language production (Figure S6)

These tasks generated three broad language-motor-attentional states (states A, B, and C), only one of them (state B) being more coupled during the behaviorally more difficult verb generation condition than during picture naming. State D corresponded mainly to a DMN state exhibiting phase desynchronization from rest to task, similarly to the *n*-Back state C.

#### S3. State classification accuracy and robust estimation of number of patterns (Figure S8)

To assess whether k-means clustering as used in the main text provided a good description of functional connectivity data across tasks, we plotted the fraction of residual variance (i.e., one minus the goodness-of-fit) of the state models as a function of the number *k* of states to be classified (Figure S8). These curves correspond to a version of the elbow curves as they provide a qualitative assessment of the balance between low residual variance and low complexity. The *k* value derived from the gap statistics is emphasized in Figure S8. The residual variance for time-averaged amplitude correlation was systematically low even at *k* = 1, with a moderate decrease at *k* = 2 (Figure S8a). This is in agreement with the gap statistics selecting either *k* = 1 or *k* = 2 depending on the dataset. The curve for time-averaged phase coupling was higher, especially at low *k* values, but presented a sharp decrease before flattening at *k* = 3 to 5 (Figure S8a). Again this agrees with the gap statistics. Finally, the residual variance for amplitude-based phase coupling (Figure S8b) and the temporal standard deviation of both short-time amplitude and phase coupling (Figure S8c, used in Results S5) were very low already at *k* = 1, which is once again consistent with the gap statistics.

Because using the right number *k* of clusters is crucial for a valid classification, we double checked the results of the gap statistics by developing a non-parametric test on the curves of residual variance shown in Figure S8. Our method relies on the observation that *k*-means clustering is an example of linear model, and is actually applicable to any categorical linear model. We identified the *k* values disclosing a statistically significant drop in residual variance when allowing one additional state in the model (i.e., when passing from *k* to *k* + 1). We then selected the first value of *k* failing to show a significant drop, which corresponds to the least complex state model for which increasing complexity does not increase the goodness-of-fit significantly. Significant drops in residual variance were detected using a minimum statistic^46,47^ on the discrete derivative of the curves shown in Figure S8. Samples of this statistic were generated non-parametrically under the null hypothesis that these curves are flat, i.e., residual variance is assumed equivalent across all *k* values. Specifically, we started from single-subject connectivity k-state models (Methods, task-dependent connectivity state classification) and randomly shuffled their *k* label (e.g., the *k* = 3 model would be seen as a *k* = 7 model, which is allowed under the null of exchangeable *k* values), independently for each subject. The minimum derivative of the resulting curve of residual variance then provided a null sample of the statistic. We used this procedure to simulate 2000 samples. Significant drops in residual variance were identified as those beneath the 5^th^ percentile of this set of null samples. In particular, the case *k* = 1 corresponded to the complete absence of any significant drop.

**Figure S2:**
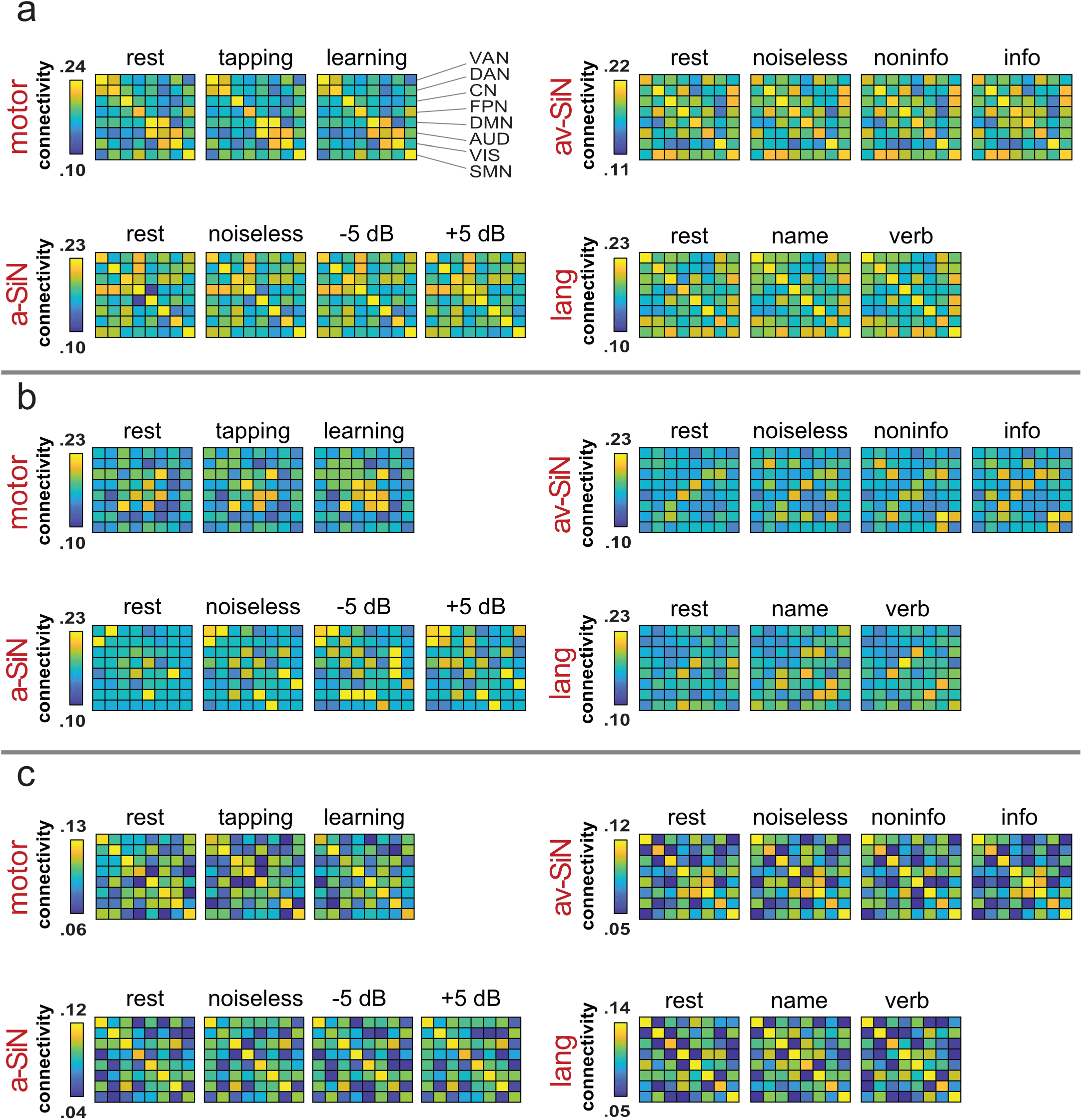
Broadband network-level connectivity. These plots show the broadband (i.e., average across frequency bands), network-level (i.e., average across RSN nodes) connectivity patterns throughout the conditions of four experiments. **a**, Amplitude correlation. **b**, Phase coupling. **c**, Amplitude-based phase coupling. motor, motor sequence learning; tapping, simple finger tapping task; learning, complex sequence learning task; a-SiN, auditory speech-in-noise comprehension; noiseless, speech comprehension task without noise; −5 dB, with informational noise at −5 dB; 5 dB, with informational noise at 5 dB; av-SiN, audiovisual speech-in-noise comprehension; noiseless, speech comprehension task without noise; noninfo, with non-informational noise; info, with informational noise; lang, covert language production; name, picture naming task; verb, verb generation task; VAN, ventral attentional network; DAN, dorsal attentional network; CN, control-executive network; FPN, fronto-parietal network; DMN, default-mode network; AUD, auditory network; VIS, visual network; SMN, sensorimotor network.

**Figure S3:**
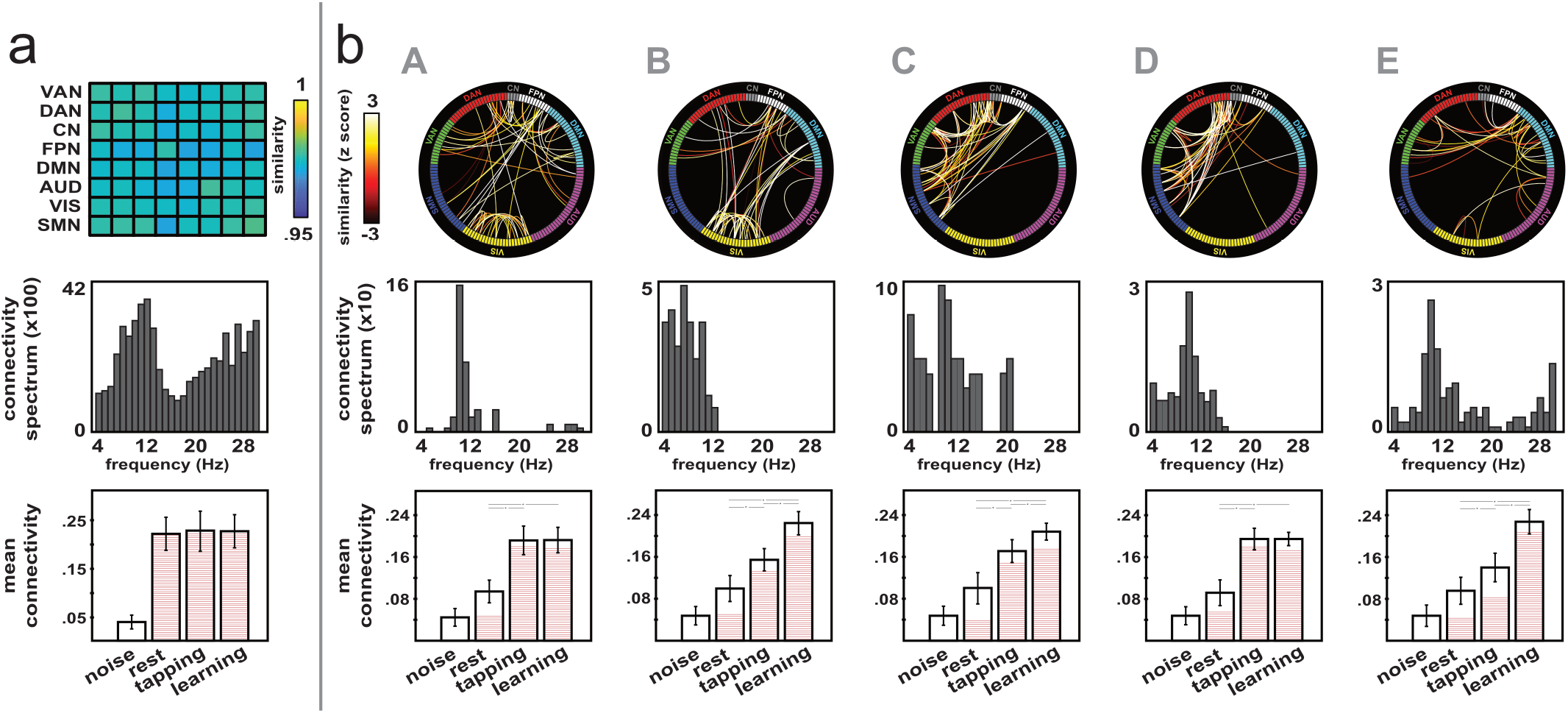
Connectivity states for the motor sequence learning experiment. **a**, Amplitude correlation. **b**, Phase coupling. Each column corresponds to one state, for which we show different characteristics: **top**, connectivity map; **middle**, spectral distribution (number of connections in the state per frequency bin); **bottom**, mean state connectivity (i.e., average across all couplings in the state) per condition, noise estimate included. Links in these connectivity maps were weighted by a measure of similarity between their task-dependent functional connectivity pattern and the state pattern shown at the bottom. For visualization purposes, the amplitude correlation state maps were summarized using coarse network-level matrices due to the overwhelming number of links to draw. For phase coupling states, the similarity values were converted into *z* scores to better emphasize color contrasts. The state connectivity bar plots at the bottom show the group mean and SEM across single-subject values. The filling in each bar indicates the proportion of state links that are significantly above noise level in the corresponding condition (*p* < 0.05 corrected for the false positive rate). Stars indicate significant differences across two conditions (t tests, *p* < 0.05, noise condition excluded). tapping, simple finger tapping task; learning, complex sequence learning task; VAN, ventral attentional network; DAN, dorsal attentional network; CN, control-executive network; FPN, fronto-parietal network; DMN, default-mode network; AUD, auditory network; VIS, visual network; SMN, sensorimotor network.

**Figure S4:**
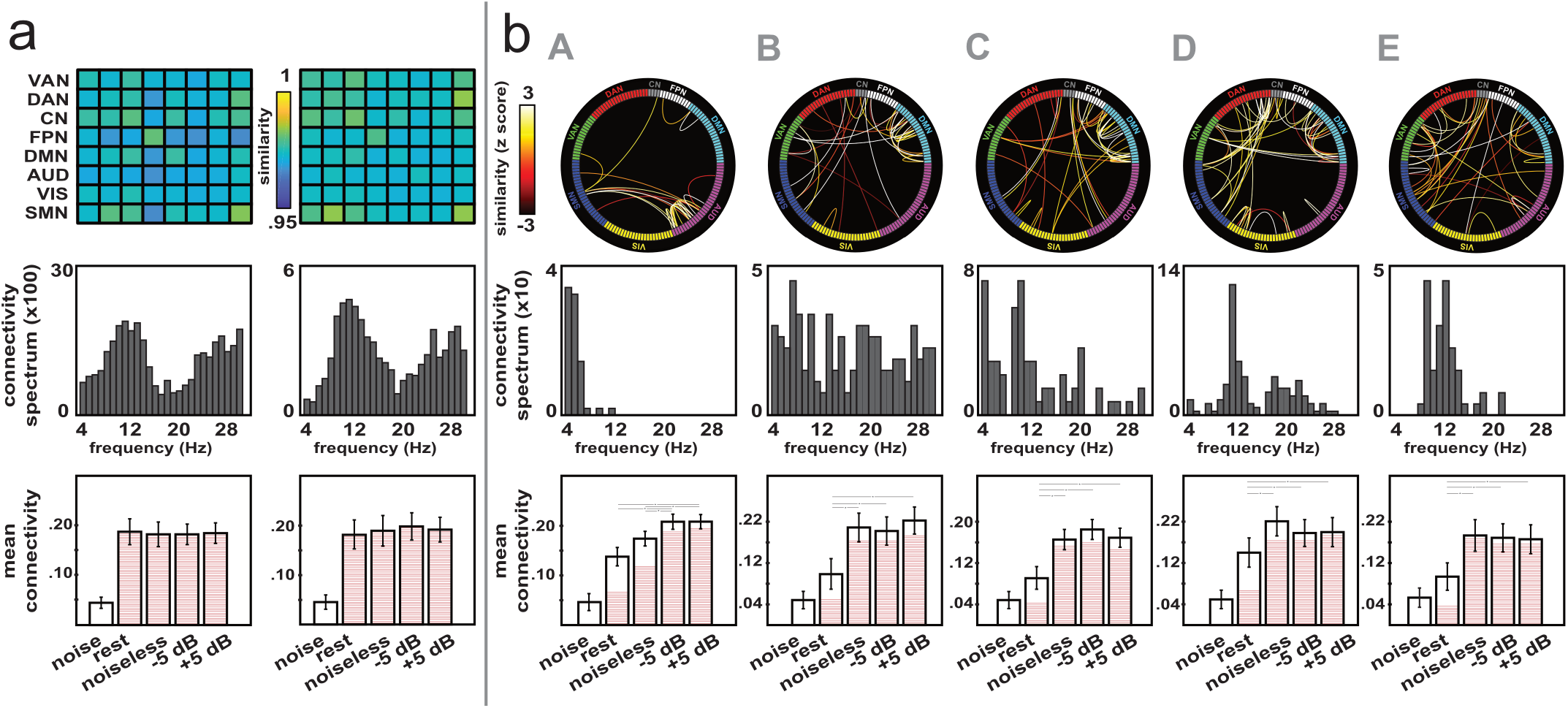
Connectivity states for the auditory speech-in-noise comprehension experiment. **a**, Amplitude correlation. **b**, Phase coupling. We refer to Figure S3 for details. noiseless, speech comprehension task without noise; −5 dB, with informational noise at −5 dB; 5 dB, with informational noise at 5 dB.

**Figure S5:**
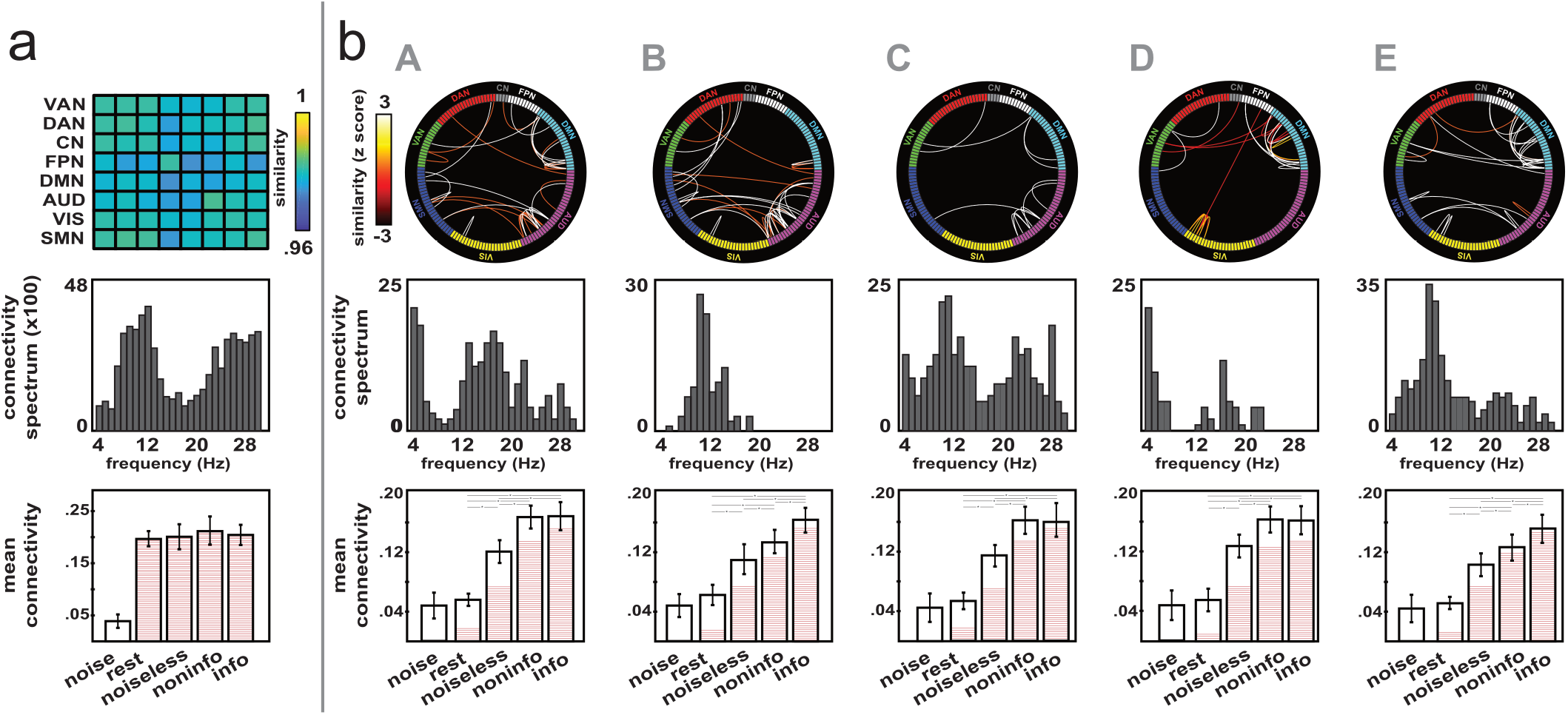
Connectivity states for the audiovisual speech-in-noise comprehension experiment. **a**, Amplitude correlation. **b**, Phase coupling. We refer to Figure S3 for details. noiseless, speech comprehension task without noise; noninfo, with non-informational noise; info, with informational noise.

**Figure S6:**
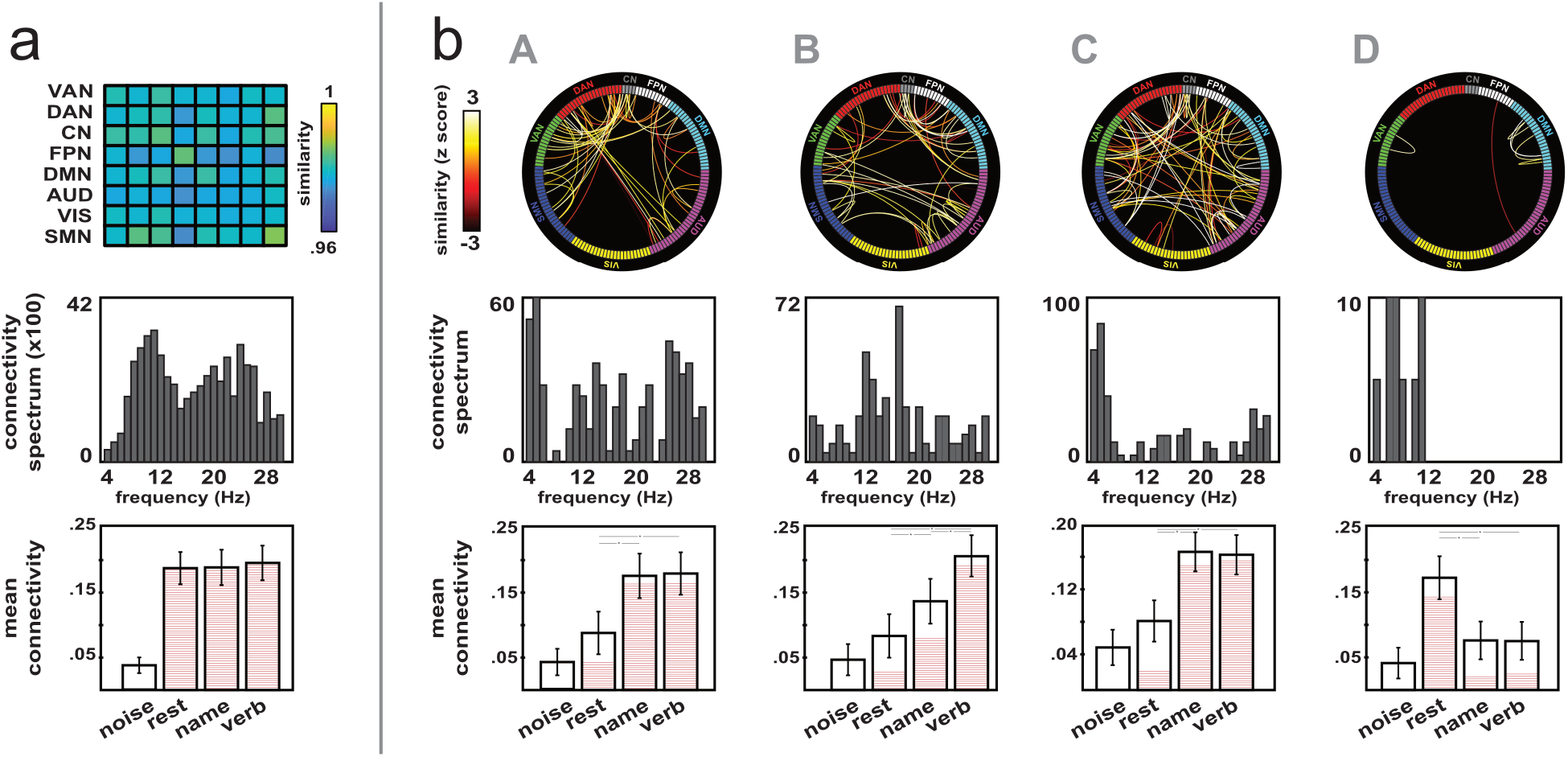
Connectivity states for the covert language production experiment. **a**, Amplitude correlation. **b**, Phase coupling. We refer to Figure S3 for details. name, picture naming task; verb, verb generation task.

**Figure S7:**
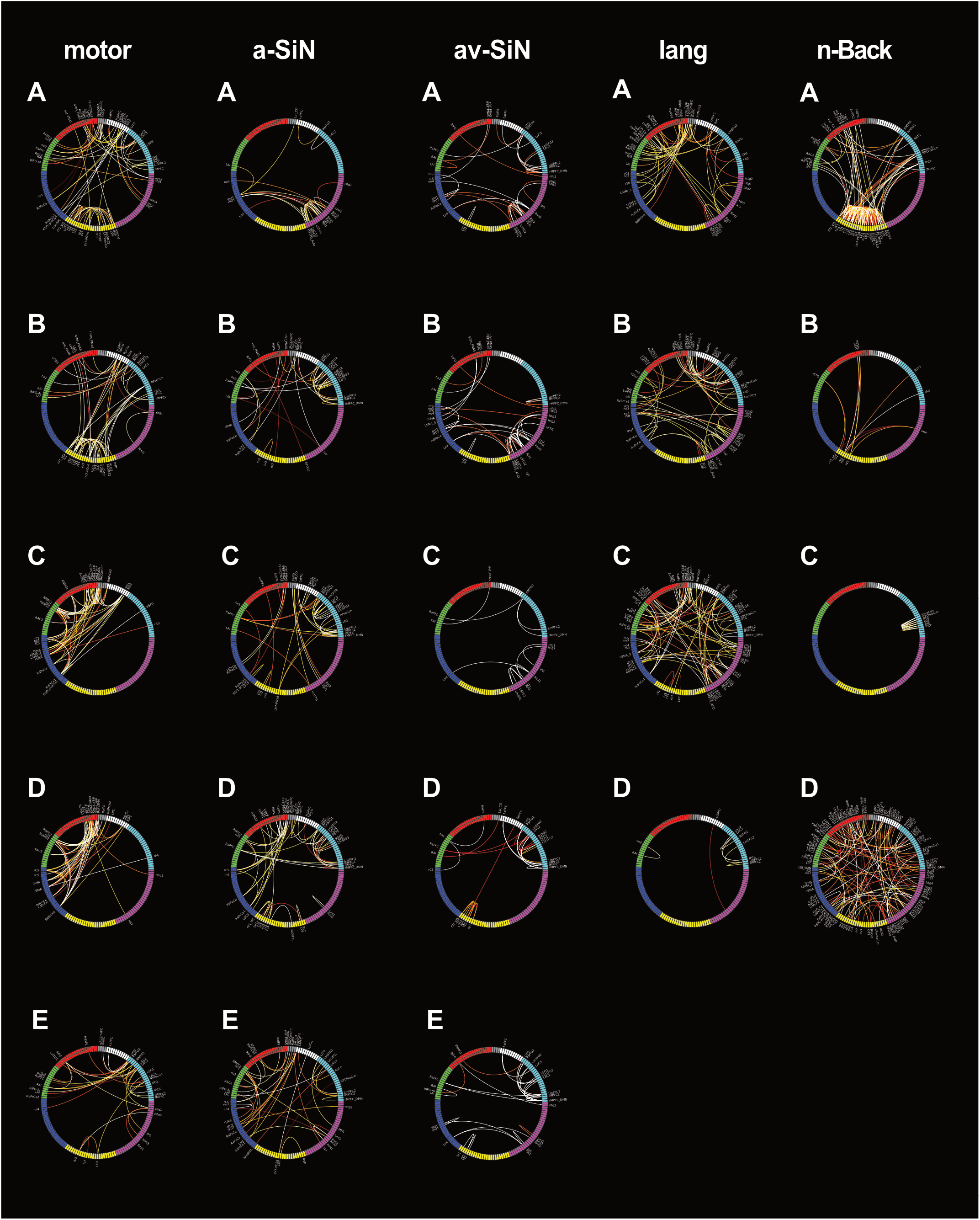
Detailed view of phase connectivity state maps. This figure gathers all phase connectivity state maps (see top part of Figures 2 and S3–S6, b), now with the label of the nodes involved in state connections. We refer to Table 1 in Della Penna et al.^40^ for the corresponding list of node labels. motor, motor sequence learning; a-SiN, auditory speech-in-noise comprehension; av-SiN, audiovisual speech-in-noise comprehension; lang, covert language production.

**Figure S8:**
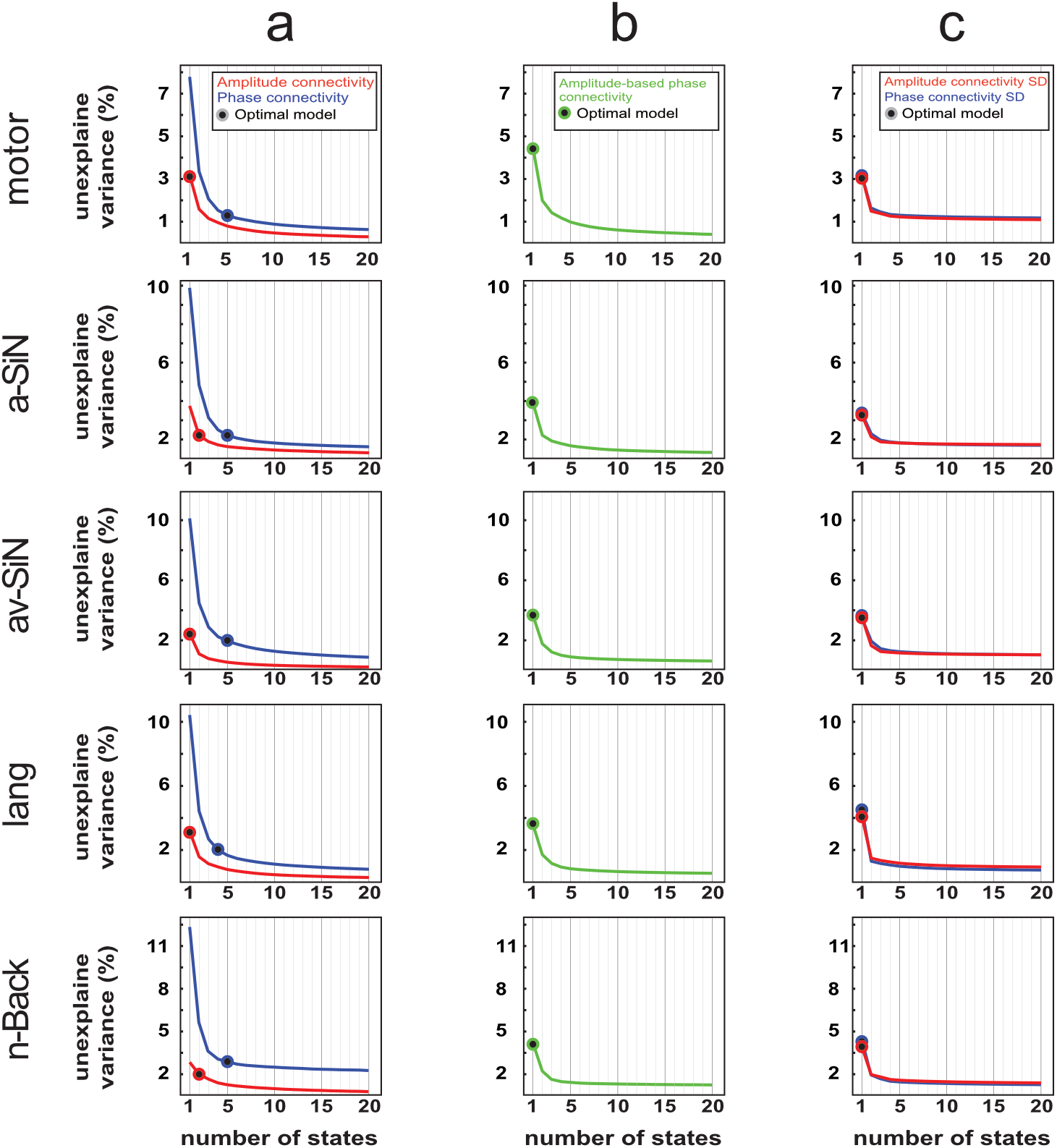
Residual variance of state classification. Elbow curves showing the fraction of residual variance for the k-means clustering models, with the number *k* of states varying between 1 and 20, for both amplitude and phase coupling. The optimal number is emphasized on these plots. **a**, Time-averaged functional connectivity. **b**, Amplitude-based model of phase connectivity. **c**, Temporal standard deviation (SD) of short-time connectivity (Results S5). motor, motor sequence learning; a-SiN, auditory speech-in-noise comprehension; av-SiN, audiovisual speech-in-noise comprehension; lang, covert language production.

Although this non-parametric approach is fairly different from the gap statistics, which relies on parametric assumptions,^50^ both approaches concurred in all the cases considered in Figure S8. This convergence thus further argues for the robustness of our state classification of task-dependent functional connectivity.

#### S4. Robustness to methodological changes (Figures S9–S14)

Our main analysis entailed methodological choices that might *a priori* affect our results. We explore here the effect of several parameters and show that they did not have a dramatic impact, using the *n*-Back dataset as case study. Only the parameter of interest was changed while keeping all the others as per the analysis reported in the main text.

##### (i) Spectral range (Figure S9)

We extended the spectral range from 4 - 30 Hz to 1 - 45 Hz (Figure S9) to include the delta (1 - 4 Hz) and low-gamma (30 - 45 Hz) bands, which are noisier but still carry functional correlates in MEG signals^16^. We did not consider frequencies beyond 45 Hz given how typically low their MEG signal-to-noise ratio is, especially with regard to RSN connectivity.^25^

##### (ii) Brain parcellation (Figure S10)

The network-based parcellation we used directly allows to assess the organization of functional connectomes into RSNs (Figure S1).^40^ To check that our results are not tied to this specific parcellation scheme, we re-computed functional connectomes with a widely-used brain atlas based on automated anatomical labelling (AAL)^52^. We placed nodes at the center-of-mass of each extended parcel and assigned them to RSNs based on their distance to the original parcellation nodes (Figure S1) to keep a uniform presentation of results.

##### (iii) Power regression (Figure S11)

Regressing out power from connectivity estimates allowed to avoid detecting possible spurious connectivity modulations due to changes in signal-to-noise ratio^45^ across experimental conditions. However, genuine connectivity modulations might also induce concurrent changes in node power,^53^ in which case this regression step might overshadow genuine task-related connectivity modulations. We checked that our results are not driven by such false negatives by repeating our analyses without the power regression step. Convergence of results would also show that they are not dramatically affected by false positives in the absence of power regression either.

##### (iv) Spatial leakage correction (Figures S12, S13)

As reviewed above (Methods, frequency-specific functional connectivity), spatial leakage correction is essential to functional connectivity mapping, yet correction methods have limitations.^32,54^ We assessed the robustness of our analysis against leakage correction by replacing our geometric correction with signal orthogonalization (Figure S12), and also by removing leakage correction altogether (Figure S13). Given that our analysis is mainly designed to detect task-related connectivity contrasts, and that spatial leakage is subdominant after contrasting (although it remains a confound^55^), we expected results to hold in both cases. Their robustness even without leakage correction would also rule out connectivity asymmetry (due to pairwise corrections^44^) as a major issue.

##### (v) Event-related fields (Figure S14)

Task-related phase connectivity changes may reflect, at least partially, event-related responses to the task at hand. These responses are by definition time locked to externally-controlled events, such as auditory or visual stimulation.^16^ Therefore, the different brain regions involved in distributed event-related activity will tend to exhibit increased phase locking. We examined to what extent this effect drove our detection of task-related phase connectivity modulations by re-estimating functional connectivity after elimination of the event-related MEG responses. We cut the MEG data into epochs from −500 ms to 1000 ms relative to each stimulus (item visual presentation identified by a trigger signal), subtracted their temporal average over the baseline (from −200 ms to 0 ms), and averaged over epochs.^14^ The contribution of the event-related response to the continuous MEG data was then modeled as the temporal convolution of this averaged epoch MEG signal with the trigger signal, and was eliminated from the continuous MEG data by subtraction. Functional connectivity was then estimated on this corrected data.

Figures S9–S14 show the task-dependent connectivity states obtained with the analyses (*i*)–(*v*) (compare to Figure 2 in the main text). State classification accurately summarized the patterns of task dependence through the *n*-Back conditions in all cases (amplitude correlation, goodness-of-fit > 93%; phase coupling, > 92%), and confirmed the task independence of amplitude correlation (F_3,102_ < 4.1, *p* > 0.26, 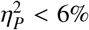 across all states and methodological variants) and the task modulations of phase coupling (F_3,102_ > 9.2, *p* < 0.002, 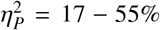). The characteristics of connectivity states remained qualitatively similar, although details differed. Either one (Figures S9, S10, S14) or two (Figures S11–S13) amplitude connectivity states emerged. In the latter cases, the two states split the spectrum into a low- and high-frequency band, as in Figure 2a, whereas in the former case the state was broadband, similarly to what was found in the other experiments (Results S2, Figures S3–S6). Each phase connectivity state described in the main text (Figure 2b) could also be identified whatever the methodological parameter changed, with similar spectrum and task-related pattern. Notably, the correction of event-related fields reduced the involvement of visual couplings in state A and displaced its connectivity spectrum from the theta band to the alpha band (compare Figure S14b with Figure 2b), but otherwise preserved all other state features. The optimal number of phase coupling states was similar (Figure S10) or higher (Figures S9, S11–S14). Consequently, in the latter cases, some states appeared split into two or three states. A notable exception was the extrinsic DMN state C exhibiting phase desynchronization during the 1- and 2-Back tasks, which remained undivided. Interestingly, the cases where possible confounds are left uncontrolled, i.e., without power regression (Figure S11) or leakage correction (Figure S13), led to substantially more phase connectivity states, which presumably reflects higher processing noise in the functional connectivity data. Still the main property of interest, i.e., their extrinsic nature, persisted.

**Figure S9:**
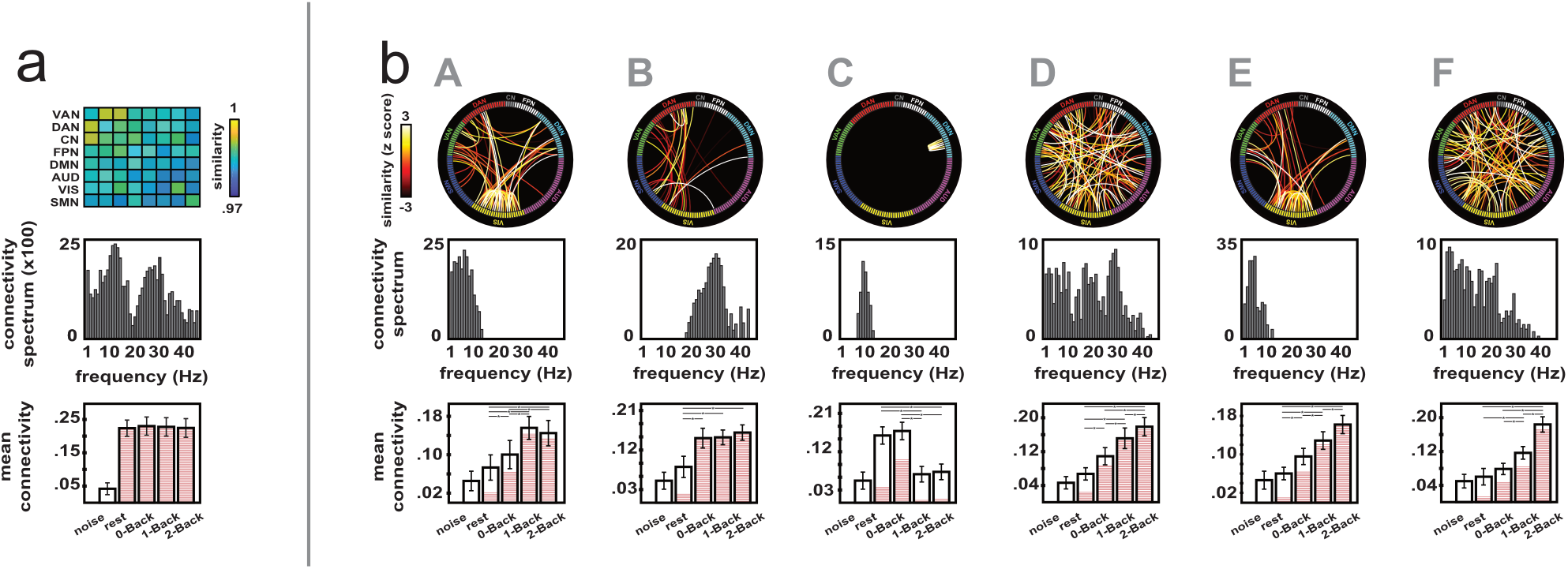
Connectivity states for the *n*-Back experiment, with wider spectral range. **a**, Amplitude correlation. **b**, Phase coupling. Similar to Figure 2, but for functional connectivity data computed over a larger frequency band (1 - 45 Hz).

**Figure S10:**
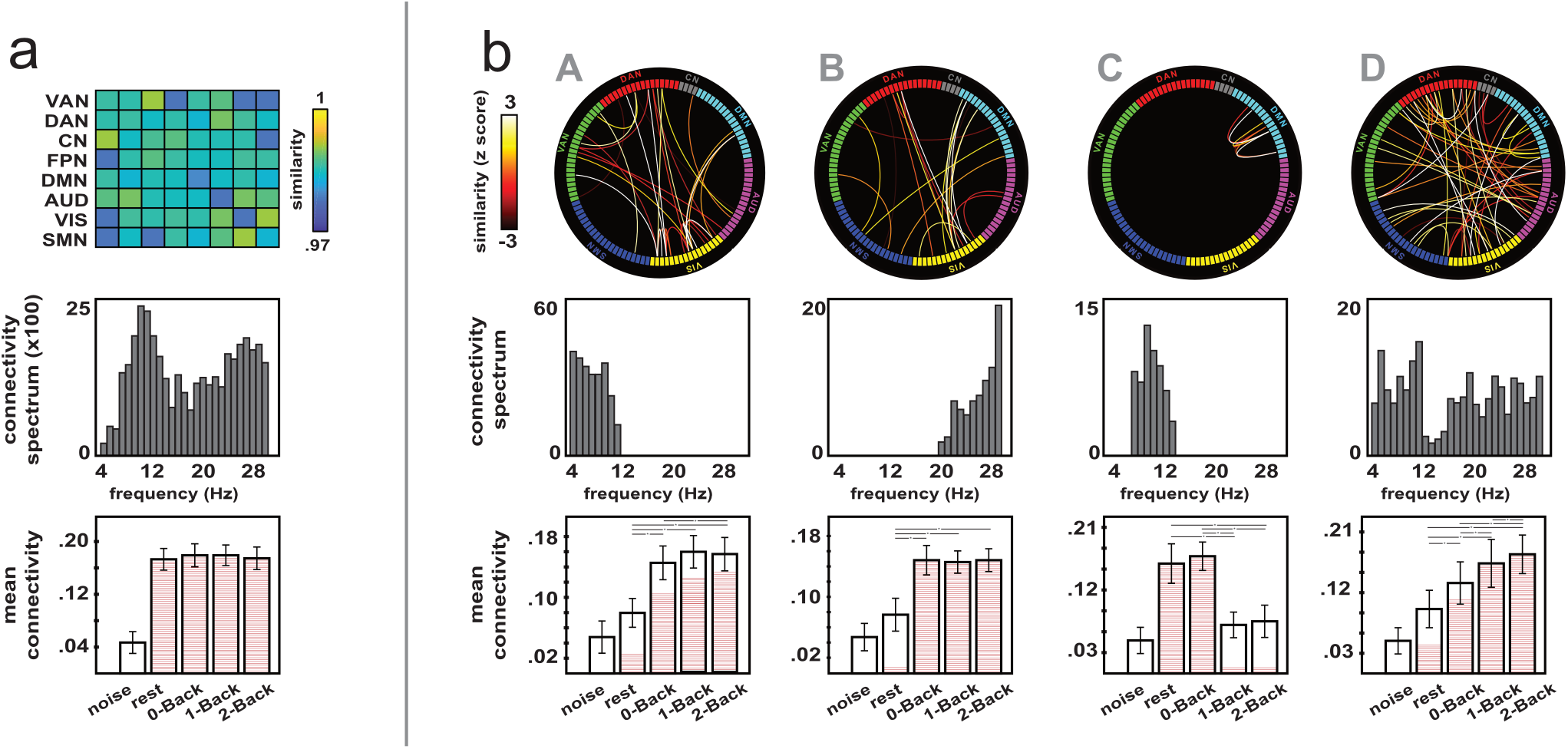
Connectivity states for the *n*-Back experiment, with another brain parcellation. **a**, Amplitude correlation. **b**, Phase coupling. Similar to Figure 2, but for functional connectivity data computed with the AAL parcellation. Network classification of AAL nodes was achieved by assigning each AAL node to the network of the closest (in Euclidean distance) node from the original brain parcellation shown in Figure S1.

**Figure S11:**
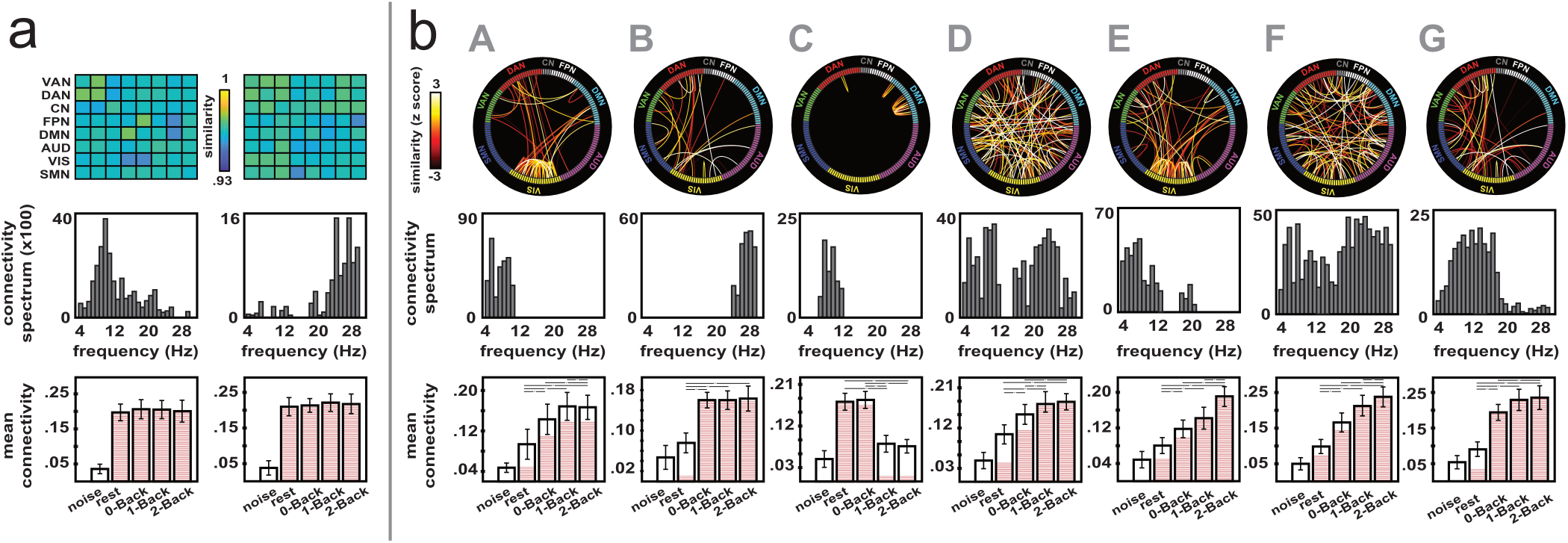
Connectivity states for the *n*-Back experiment, without power regression. **a**, Amplitude correlation. **b**, Phase coupling. Similar to Figure 2, for functional connectivity data without power regression.

**Figure S12:**
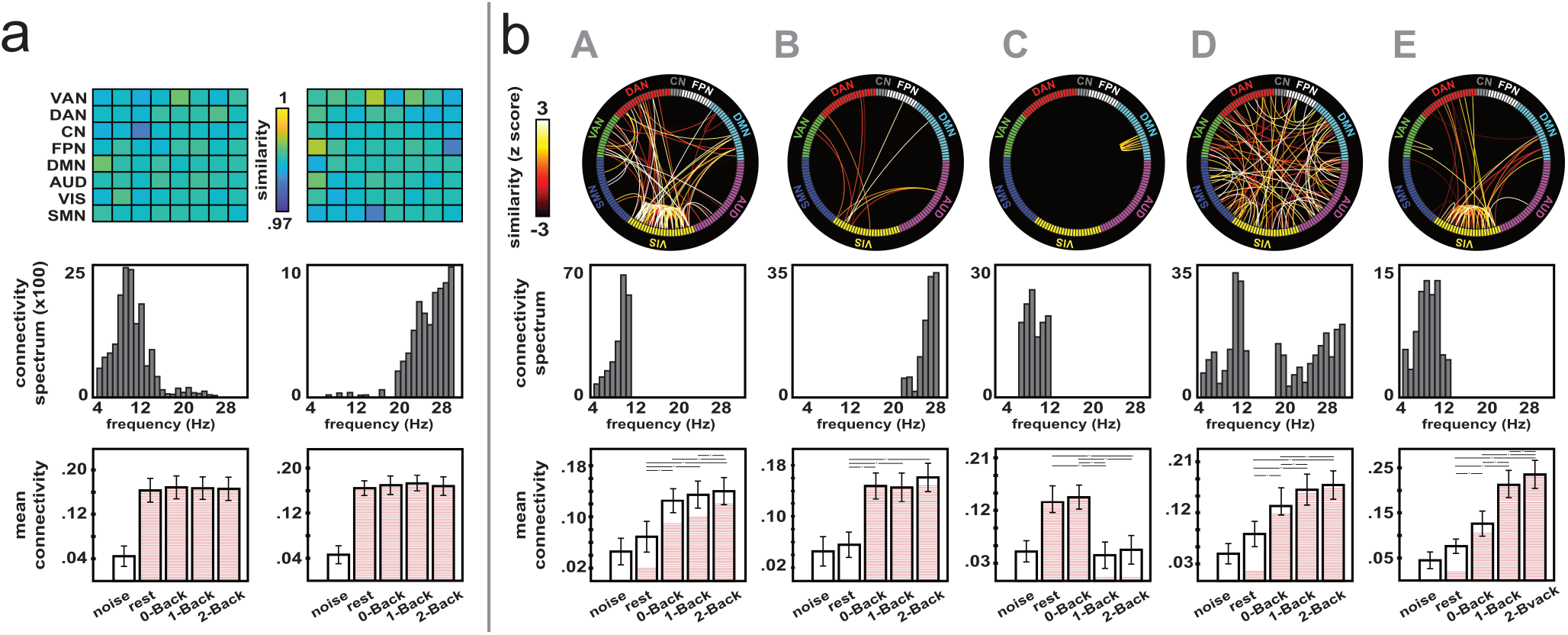
Connectivity states for the *n*-Back experiment, with signal orthogonalization. **a**, Amplitude correlation. **b**, Phase coupling. Similar to Figure 2, for functional connectivity data with spatial leakage corrected via signal orthogonalization.

**Figure S13:**
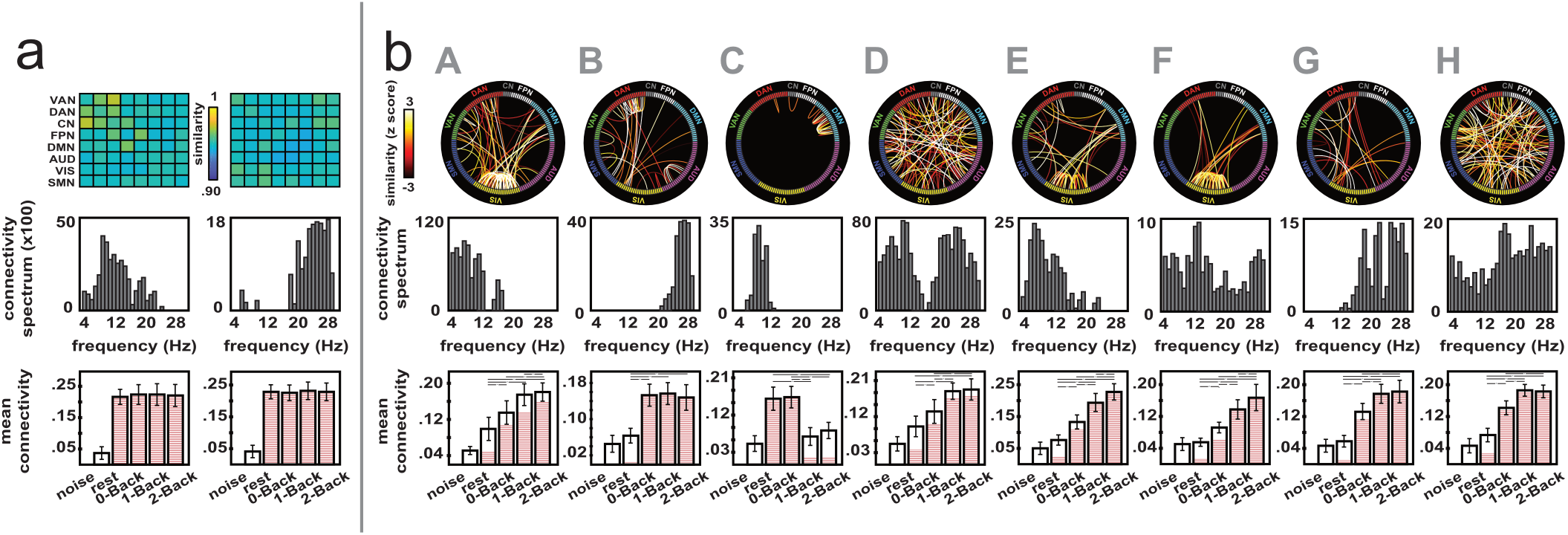
Connectivity states for the *n*-Back experiment, without leakage correction. **a**, Amplitude correlation. **b**, Phase coupling. Similar to Figure 2, for functional connectivity data with spatial leakage left uncorrected.

**Figure S14:**
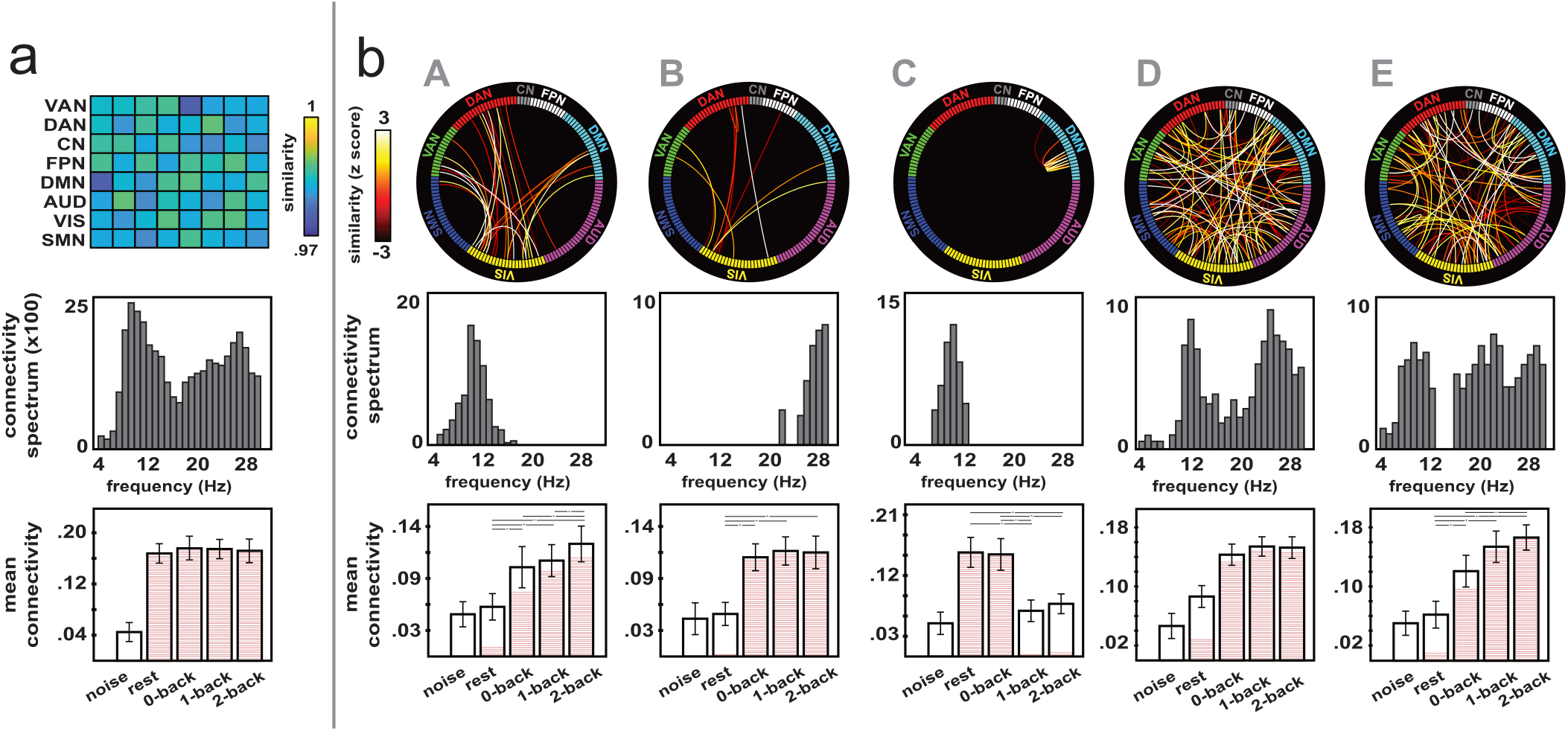
Connectivity states for the *n*-Back experiment, with correction for event-related responses. **a**, Amplitude correlation. **b**, Phase coupling. Similar to Figure 2, for functional connectivity data obtained after elimination of event-related MEG responses.

These analyses thus confirmed the robustness of our results against the methodological variations (*i*)–(*v*). The last case (*v*) specifically deserves further discussion, especially in light of the conceptual framework developed in the main text. Should we exclude task-related modulations in phase connectivity explained by event-related responses? The analog of this effect in slow-wave fMRI signals is considered to confound task-related connectivity,^56^ but at the higher temporal resolution of MEG the situation is different. Distributed event-related responses generally emerge from functional integration and thus reflect precisely task-related neural communication. The only methodological confound is that stimulation may co-activate separate regions without them actually communicating, but this possibility is restricted to zero-lag phase synchronization between homologous primary cortices. This agrees with our observation that correcting for event-related responses only removed intra-visual theta-band couplings in the *n*-Back phase connectivity state A of Figure 2b (Figure S14b). Since no task-dependent phase connectivity state was limited to primary cortices (as attentional networks were systematically involved), we conclude than they all reflected genuine functional connectivity modulations. More conceptually, event-related responses only represent a part of active, extrinsic neural processes. They are highlighted, somewhat artificially, because we control them experimentally and we can thus disambiguate them from background brain activity. However, several uncontrolled active processes will also take place, e.g., spontaneous cognition or sustained task-related attention, but not directly time locked to stimuli. Accordingly, we surmise that such processes contribute to extrinsic phase coupling and were detectable with our block-design, time-averaged connectivity analysis. The *n*-Back phase connectivity states shown in Figure S14 would indeed precisely reflect these extrinsic processes. The DMN state C provides a striking example of this statement. It exhibits high phase coupling in conditions of low cognitive load (here, rest and 0-Back), possibly reflecting active spontaneous cognitive processes such as recurrent mind-wandering episodes^57^. This state then decouples in conditions of high cognitive load (here, 1- and 2-Back) as attentional resources are shifted towards external tasks and the opportunity to shift towards spontaneous cognitive processes is drastically reduced.

#### S4. Temporal fluctuations of short-time functional connectivity (Figures S15, S16)

We claimed in the main text that the relationship determined between time-averaged amplitude and phase coupling at the group level (Figure 4a) generalizes to short-time connectivity dynamics. This result detailed here was obtained by analyzing the temporal fluctuations of sliding-window connectivity around their time average (i.e., after mean centering). We estimated the Spearman correlation between the time-locked fluctuations of amplitude and phase coupling, separately for each frequency-specific connection disclosing time-averaged phase locking significantly above noise level, and for each experimental condition. Connectivity time series were temporally concatenated across subjects beforehand to generate a group-level analysis. Importantly, the inter-subject correlation effect reported in Figure 4a was excluded thanks to mean centering, so this analysis specifically focused on the intra-subject, dynamic effect. The distribution of these correlations was clearly shifted towards positive values (Figure S15), with a significantly positive mean (p < 0.001 across conditions and datasets). Accordingly, 42 - 56% of correlations (range across conditions and datasets) were significantly positive at *p* < 0.05 corrected for multiple comparisons (Methods, cross-correlation of amplitude and phase connectivity, for a description of these statistical procedures.) We used this result in the main text to justify our construction of the amplitudebased model of phase coupling.

**Figure S15:**
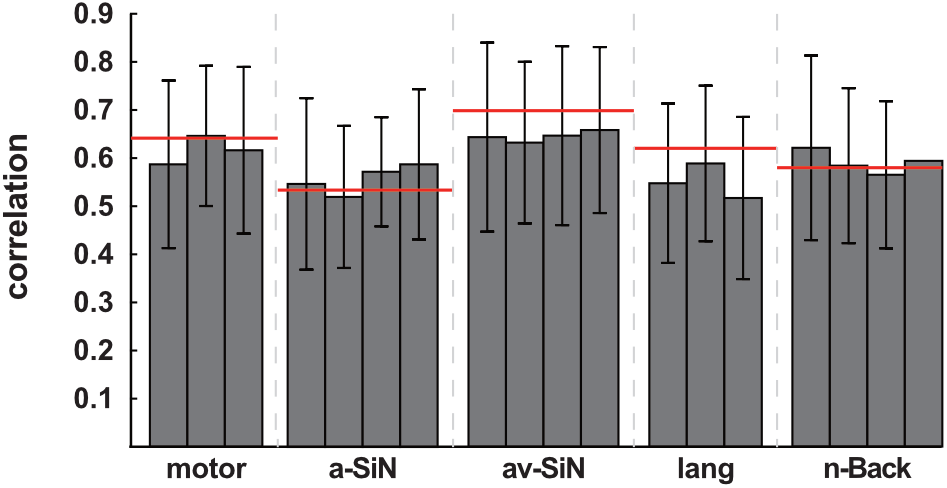
Temporal correlation between short-time amplitude and phase coupling fluctuations. Mean and SD of temporal Spearman correlations between short-time fluctuations of amplitude and phase connectivity in each condition of our five experiments. Each bar corresponds to one condition. Only the connections that were included in time-averaged phase connectivity state classification contributed to this distribution. Thresholds for correlation significance (*p* < 0.05 corrected for the false positive rate) are indicated in red. motor, motor sequence learning; a-SiN, auditory speech-in-noise comprehension; av-SiN, audiovisual speech-in-noise comprehension; lang, covert language production.

What this result also entails is that short-time phase coupling fluctuations may be expected to exhibit intrinsic dynamics closely tied to that of amplitude correlation. To further illustrate this claim, we describe here an analysis of the temporal standard deviation (SD) of short-time couplings (rather than their time average as done in the main text). First, the whole connectome (i.e., all connections and frequencies) revealed that both amplitude and phase sliding-window connectivity SD was significantly higher than their noise estimates over the entire connectome (p < 0.05 with false positive rate controlled by maximum statistics, along the lines described in Methods, frequency-specific functional connectivity). This simple yet crucial observation demonstrates the existence of a short-time brain coupling dynamics beyond mere statistical variability (a point sometimes debated in the literature^43,58^). Second, while time-averaged amplitude correlation was stronger within RSNs (Figures 1c and S2a), amplitude correlation SD emerged most prominently between RSNs. This shows that cross-RSN integration is temporally more unstable than within-RSN integration (Figure S16a). This is in line with data putting cross-RSN amplitude coupling at the center of dynamic brain integration^36,43,59,60^. Phase-locking SD did not appear structured that way (Figure S16b), which is reminiscent of its time average (Figures 1d and S2b). Finally, we examined whether connectivity SD was modulated by task using *k*-means clustering (as was used on time-averaged connectivity in the main text). Both amplitude and phase coupling required a single state of connectivity SD (goodness-of-fit > 96% across datasets and coupling types; Figure S8c), which was task independent (ANOVA; *F* < 3.9, *p* > 0.34, 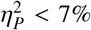; Figures S16c–e). We conclude that the temporal variability of both short-time amplitude and phase coupling is dominated by an intrinsic dynamics. Our results in the main text show that this intrinsic dynamics becomes subdominant after time averaging over a few minutes.

**Figure S16:**
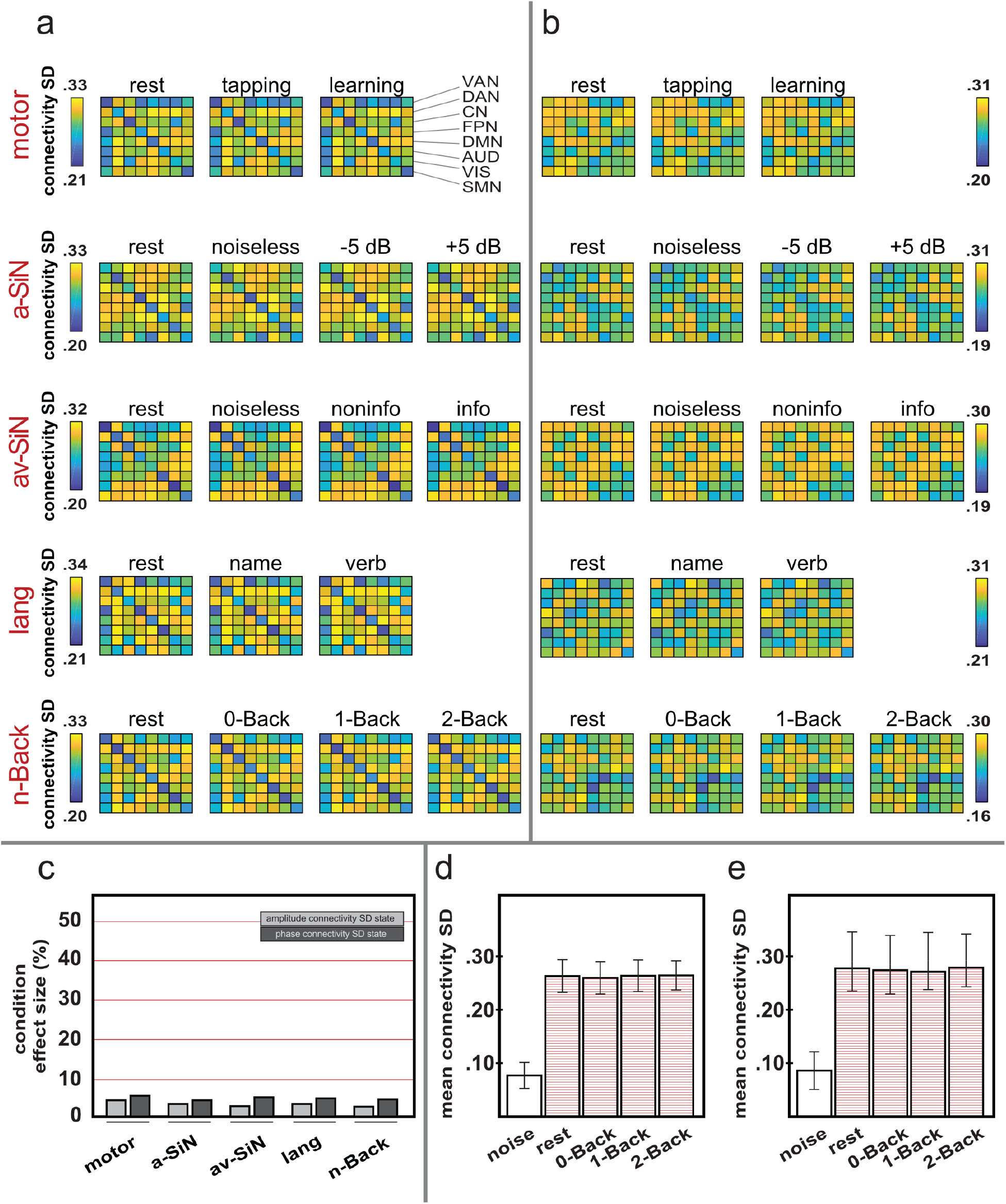
Temporal variability in short-time connectivity fluctuations. The spatial patterns of connectivity SD are illustrated using broadband (i.e., average across frequency bands), network-level (i.e., average across RSN nodes) plots. **a**, Amplitude correlation. **b**, Phase coupling. Part **c** shows the effect sizes of the ANOVA applied to task-dependent connectivity SD averaged across all connections and frequencies, for each experiment (light grey, amplitude correlation; dark grey, phase coupling). These small effect sizes are illustrated in the case of the *n*-Back experiment by plotting the mean connectivity SD (i.e., average across all connections and frequencies) as a function of condition (noise estimate included). **d**, Amplitude correlation. **e**, Phase coupling. motor, motor sequence learning; tapping, simple finger tapping task; learning, complex sequence learning task; a-SiN, auditory speech-in-noise comprehension; noiseless, speech comprehension task without noise; −5 dB, with informational noise at −5 dB; 5 dB, with informational noise at 5 dB; av-SiN, audiovisual speech-in-noise comprehension; noiseless, speech comprehension task without noise; noninfo, with non-informational noise; info, with informational noise; lang, covert language production; name, picture naming task; verb, verb generation task; VAN, ventral attentional network; DAN, dorsal attentional network; CN, control-executive network; FPN, fronto-parietal network; DMN, default-mode network; AUD, auditory network; VIS, visual network; SMN, sensorimotor network.

